# Complement C3 and C3aR mediate different aspects of emotional behaviours; relevance to risk for psychiatric disorder

**DOI:** 10.1101/685537

**Authors:** Laura J. Westacott, Trevor Humby, Niels Haan, Sophie A. Brain, Emma-Louise Bush, Margarita Toneva, Andreea-Ingrid Baloc, Anna L. Moon, Jack Reddaway, Michael J. Owen, Jeremy Hall, Timothy R. Hughes, B. Paul Morgan, William P. Gray, Lawrence S. Wilkinson

## Abstract

Complement is a key component of the immune system with roles in inflammation and host-defence. Here we reveal novel functions of complement pathways impacting on emotional reactivity of potential relevance to the emerging links between complement and risk for psychiatric disorder. We used mouse models to assess the effects of manipulating components of the complement system on emotionality. Mice lacking the complement C3a Receptor (*C3aR*^-/-^) demonstrated a selective increase in unconditioned (innate) anxiety whilst mice deficient in the central complement component C3 (*C3*^-/-^) showed a selective increase in conditioned (learned) fear. The dissociable behavioural phenotypes were linked to different signalling mechanisms. Effects on innate anxiety were independent of C3a, the canonical ligand for C3aR, consistent with the existence of an alternative ligand mediating innate anxiety, whereas effects on learned fear were due to loss of iC3b/CR3 signalling. Our findings show that specific elements of the complement system and associated signalling pathways contribute differentially to heightened states of anxiety and fear commonly seen in psychopathology.

## 1. Introduction

The complement system is a key component of the immune system that plays a pivotal role in inflammation and host-defence. Complement activation occurs via several pathways, all of which lead to cleavage of the central protein, C3 (see Figure S1). Activation of C3 generates the fragments C3a and C3b. C3a is an anaphylatoxin that signals via its canonical G-protein coupled receptor C3aR^1^. Activation of this receptor has been demonstrated to trigger calcium mobilization^2-4^, stimulating an array of intracellular signalling pathways to induce both pro- and anti-inflammatory effects^1,5^. C3b on the other hand propagates further complement activation by contributing to the cleavage of complement component 5 (C5) downstream of C3 and, after further cleavage to iC3b, plays a role in opsonisation by macrophages and microglia via complement receptor 3 (CR3). Akin to C3, C5 cleavage generates C5a (another anaphylatoxin and a ligand for the C5a receptor, C5aR) and C5b, which triggers the terminal complement pathway by sequentially binding proteins C6, C7, C8 and C9. These proteins subsequently congregate to assemble the membrane attack complex (MAC) which ultimately results in destruction of the target cell or pathogen via cell lysis^6^.

In the central nervous system evidence is emerging that complement has functions beyond its canonical immune roles^7^. Neurons, astrocytes and microglia express complement receptors and regulators, and are also capable of synthesising complement proteins^8,9^. The expression patterns of these vary over the course of brain development^10^. Complement impacts a number of neurodevelopmental processes including neurogenesis^11^, migration^12^ and synaptic elimination^13^ as well as ongoing synaptic plasticity processes underlying learning and memory in the adult brain^14^.

Furthermore, there is increasing evidence that complement is causally involved in the pathogenesis of neurodegenerative and psychiatric conditions. In Alzheimer’s disease, genetic variants in complement related loci have been associated with increased disease risk^15,16^, and complement knockout mice exhibit reduced age-related synapse loss^17^ and neuropathology^18^. Alterations in complement proteins and activation have also been reported in sera from individuals with autism-spectrum disorder^19^ schizophrenia^20^, major depressive disorder^21^, bipolar disorder^22^ and post-traumatic stress disorder^23^. In the case of schizophrenia, an important finding comes from elegant genetic work demonstrating that structural variation in the complement *C4A* locus is associated with risk of developing the disease^24^. C4 cleavage generates fragments that contribute to the activation of C3, yielding C3a and C3b. Given the known roles for the iC3b/CR3 pathway in developmental synaptic pruning^13,25^, it has been suggested that *C4A* variants may impact on psychiatric risk via this mechanism, with excessive synaptic elimination leading to abnormal connectivity and disruption of neural networks^24^. Variants in *C3* and putative complement-control genes *CSMD1* and *CSMD2* have also been implicated in genetic susceptibility for schizophrenia^26,27^.

Altered emotional function, in particular maladaptive anxiety and fear, is a pervasive and clinically important symptom in schizophrenia and a frequent comorbidity across several of the DSM-5 and ICD-11 defined disorders. Anxiety and fear exist along a spectrum of aversive emotional states and can be elicited by differing environmental factors to result in distinguishable behavioural outputs^28^. Anxiety is characterised by sustained arousal, hypervigilance and risk assessment surrounding anticipated or potential threats, while fear is often characterised as an acute response to an experienced, imminent danger resulting in immediate avoidance, fight or freezing behaviour^29,30^. Whilst there is significant overlap in the neurocircuitry underlying these states, there are also contributions from distinct neuronal circuitries^28,31^.

There is previous data suggesting complement may play a role in emotional responses to aversive circumstances. Mice overexpressing the human *C4A* variant associated with risk for schizophrenia demonstrated elevated anxiety behaviour^32^. Anxiety phenotypes have also been reported in mice exposed to excessive pre-natal complement activity^33^ and neurodegeneration-associated anxiety phenotypes are reduced by complement inhibitors^34^. Furthermore, aged *C3* deficient mice exhibited lower levels of anxiety alongside enhanced learned fear responses^17^, whereas increased anxiety has been reported in mice lacking the C3aR^35^. These previous studies suggest that complement can influence both innate and learned aversive behaviours, however, the precise complement signaling pathways responsible for effects on these dissociable aspects of emotionality is unknown.

Utilising the central role of C3 in complement signalling, we used a combination of complement knockout mice to functionally parse innate anxiety and learned fear related phenotypes. In homozygous *C3* knockout mice (*C3*^-/-^)^36^ complement cannot be activated beyond C3, and therefore these animals lack C3 activation fragments (C3a, C3b) and downstream activation products (C5a, C5b) and thus cannot activate the terminal complement pathway. Phenotypes in this model could therefore be the result of loss of any of these downstream effector molecules. We compared the *C3*^*-/-*^ model with homozygous *C3aR* knockout mice (*C3aR*^-/-^)^37^. In these mice, complement is intact apart from the capacity for C3a to bind its canonical receptor C3aR and hence through use of both models, we tested the extent to which phenotypic effects were the result specifically of disrupted C3a/C3aR signalling. A priori, because C3a is an obligate cleavage fragment of C3, we hypothesised that any phenotypes dependent on interaction of C3a and C3aR would be apparent in both *C3*^-/-^ and *C3aR*^-/-^ models.

Contrary to our initial hypotheses we found clear phenotypic dissociations between the *C3*^-/-^ and *C3aR*^-/-^ models dependent on the nature of the emotional behaviour. *C3aR*^-/-^ mice displayed a profound innate anxiety phenotype that was lacking in the *C3*^-/-^ model. In contrast, when we examined learned fear, where a previously neutral cue generates a fear response as a result of predicting an aversive outcome, we found that the dissociation was reversed with the *C3*^-/-^ mice exhibiting an enhanced fear response to a conditioned cue, but no differences between wildtype and *C3aR*^-/-^mice. This double dissociation in emotionality was linked to distinct underlying complement signalling mechanisms, with anxiety phenotypes in C3aR mice likely to be independent of C3a/C3aR signalling raising the possibility of an alternative ligand for C3aR, and *C3*^-/-^ effects on learned fear likely the result of alterations in iC3b-CR3 pathway activity. These findings point to a hitherto unrecognized complexity of complement effects on brain function and behaviour of relevance to emotional dysfunction in psychopathology.

## 2. Materials and Methods

### 2.1 Mouse models and husbandry

Wildtype and *C3*^-/-^ strains were sourced in-house from Professor B. Paul Morgan and Dr Timothy Hughes (strains originally from The Jackson Laboratory; B6.PL-Thy1^a^/CyJ stock#000406 and B6;129S4-C3tm1Crr/J stock#003641 respectively); *C3aR*^-/-^ mice were provided by Professor Craig Gerard of Boston Children’s Hospital, USA (strain subsequently provided to The Jackson Laboratory; B6.129S4(C)-*C3ar1*_*tm1Cge*_ /BalouJ; stock#033904). *C5* ^-/-^ mice (as described in ^38^) were provided by Professor Marina Botto, Imperial College London. This strain originated from naturally C5-deficient DBA/2J mice, that had been backcrossed to C57Bl/6J. *C3*^-/-^, *C3aR*^-/-^ and *C5*^-/-^ strains were maintained via homozygous × homozygous breeding and were on a C57Bl/6J background. In all experiments, knockout mice were compared to wildtype mice also on a C57Bl/6J background. Mice were between 3-8 months old during experimental testing and were kept in a temperature and humidity-controlled vivarium (21±2°C and 50±10%, respectively) with a 12-hour light-dark cycle (lights on at 07:00hrs/lights off at 19:00hrs). Home cages were environmentally enriched with cardboard tubes, soft wood blocks and nesting materials and animals were housed in single sex littermate groups (2-5 mice/cage). Standard laboratory chow and water were available *ad libitum*. All procedures were performed in accordance with the requirements of the UK Animals (Scientific Procedures) Act (1986).

### 2.2 General behavioural methods

Testing took place between the hours of 09:00 and 17:00, with random distribution of testing for subjects of different genotypes throughout the day. Mice were habituated to the test rooms for 30 min prior to testing. All assays involved individual testing of mice and apparatus was cleaned thoroughly with a 70% ethanol solution between subjects.

### 2.3 Data collection

Data for the EPM, EZM and Open Field were collected using EthoVision XT software (Noldus Information Technology, Netherlands) via a video camera mounted above the centre of each piece of apparatus. Tracking of each subject was determined as the location of the greater body-proportion (12 frames/s) in the specific virtual zones of each piece of apparatus.

### 2.4 The elevated plus maze (EPM)

The maze, positioned 300 mm above the floor and illuminated evenly at 15 lux, was constructed of opaque white Perspex and consisted of two exposed open arms (175 × 78 mm^2^, length × width, no ledges) and two equally sized enclosed arms, which had 150 mm high walls^39^. Equivalent arms were arranged opposite one another. Subjects were placed at the enclosed end of a closed arm and allowed to freely explore for 5 minutes. Data from each pair of arms were combined to generate single open and closed arm values (number and duration of arm entries and latency of first entry to each arm). In addition, the following parameters were manually scored (by an experimenter positioned at a computer in the same room as the maze, watching the live-video stream of the test); number of stretch-attend postures (SAPs; defined as the animal slowly and carefully reaching towards the open arms in a low, elongated body posture^40,41^) and number of head dips from the open arms (looking down over the edge of an open arm).

### 2.5 The elevated zero maze (EZM)

The maze, positioned 520 mm above the floor and illuminated evenly at 15 lux, was constructed of wood and consisted of two exposed open regions (without ledges; 52 mm wide) and two equally sized enclosed regions (also 52 mm wide), which had 200 mm high grey opaque walls. The diameter of the maze was 600mm. Equivalent regions were arranged opposite one another. Subjects were placed at the border of one of the open and closed regions and allowed to freely explore for 5 min. Data from each pair of regions were combined to generate single open and closed region values (number and duration of region entries and latency of first entry to each region). In addition, the number of head dips (as above) were measured. Due to the high walls of the enclosed sections of the maze, subjects were not visible to the experimenter when in the closed regions and therefore these parameters were scored only when a subject was on the open regions.

### 2.6 Locomotor activity (LMA)

LMA was measured in an apparatus consisting of twelve transparent Perspex chambers (each 210 × 360 × 200 mm, width × length × height). Two infrared beams were embedded within the walls of each chamber, which crossed the chamber 30 mm from each end and 10 mm from the chamber floor. Individual subjects were placed in a designated chamber for a 120 min duration on three consecutive days. Beam breaks were recorded as an index of activity, using a computer running custom written BBC Basic V6 programme with additional interfacing by ARACHNID (Cambridge Cognition Ltd, Cambridge, UK). Data were analysed as the total number of beam breaks per session per day.

### 2.7 Fear-potentiated startle (FPS)

FPS was assessed using startle chamber apparatus which consisted of a pair of ventilated and soundproofed SR-LAB startle chambers (San Diego Instruments, CA, USA) each containing a non-restrictive Plexiglas cylinder (35 mm in diameter), mounted on a Perspex plinth, into which a subject was placed. The motor responses of subjects to white noise stimuli (generated from a speaker 120 mm above the cylinder) were recorded via a piezoelectric accelerometer, attached centrally below the Plexiglas cylinder, which converted flexion plinth vibration into electrical signals. The peak startle response, within 200ms from the onset of each startle presentation, in each trial, was normalized for body weight differences using Kleiber’s 0.75 mass exponent^42^ as per^43^. A computer running SR-Lab software (Version 94.1.7.48) was used to programme trials and record data. A foot shock grid connected to a shock generator (San Diego Instruments, CA, USA) was inserted into the Plexiglas cylinder before conditioning sessions.

FPS consisted of three separate sessions presented over a two-day period (see Figure 4A). On the first day, mice were given a pre-conditioning session immediately followed by the conditioning session. The pre-conditioning session started with a 5 min acclimatisation phase followed by presentation of 3 no-stimulus trials, and then a block of pulse-alone trials presented at 90, 100 and 110dB (5 of each at 40 ms duration). Trials were randomly distributed throughout the session and presented with a 60 s random interval (range 36 s to 88 s). After the pre-conditioning session was complete, mice were removed from the startle chambers, restraint tubes cleaned, and shock grids were placed into the Plexiglas cylinders prior to commencing the conditioning session. The mice were then returned to the startle chambers and subjected to a session consisting of a 5 min acclimatisation phase followed by 3 CS+shock trials, with 3 no stimulus trials before and after, presented with a 2min random interval (range 1.5 to 3min). The scrambled 0.14 mA, 0.5 s foot shock was delivered in the final 0.5 s of the 30 s visual CS. Following a 24hr delay, subjects were assessed for FPS in the post-conditioning session. This session followed the same format as the pre-conditioning session (5 min acclimatisation phase followed by presentation of 3 no-stimulus trials, and then a block of pulse-alone trials presented at 90, 100 and 110dB, with 5 of each at 40 ms duration) however the final block of trials also included pulse+CS trials at 90, 100 and 110 dB (5 of each), with the startle pulse presented in the final 40 ms of the CS. FPS was determined as the fold change between pulse-alone trials and pulse+CS trials within the post-conditioning session.

### 2.8 Corticosterone measurements

Testing took place between the hours of 10:00 and 14:00 to account for the diurnal pattern of corticosterone release^44^. Mice were allowed to freely explore the EPM for 5 min, after which they were placed in a holding cage for a further 25 min before being culled by cervical dislocation. Control mice were removed from their home cage and immediately culled. There was an equal distribution of subjects of different genotypes, counterbalanced between the two test conditions and throughout the testing period. Trunk blood was collected into heparin tubes (Becton Dickinson, USA) and immediately centrifuged at 4000 rpm for 10 min, and the supernatant removed and frozen at -80°c until further use. A corticosterone ELISA was performed according to manufacturer’s instructions (ADI-900-097, Enzo Life Sciences, UK) and analysed using a four-parameter logistic curve plug in (https://www.myassays.com/four-parameter-logistic-curve.assay).

### 2.9 Diazepam study

Wildtype, *C3*^*-/-*^ and *C3aR*^*-/-*^ were used and were randomly assigned to either vehicle or drug conditions within each genotype. A three-day dosing regimen of diazepam (2 mg/kg, i.p., Hameln Pharmaceuticals, UK) or an equivalent volume of vehicle (0.1 M phosphate buffered saline, pH 7.4) was used, based on pilot testing in wildtype mice to establish an effective anxiolytic dose with minimal sedative effects (data not included). Following 2 days of pre-treatment, diazepam or vehicle was administered 30 min prior to testing on the EPM on the 3^rd^ day.

### 2.10 Tissue for gene expression analysis

Mice were removed from their home cage and immediately culled via cervical dislocation. Brains were removed and the following regions dissected: medial prefrontal cortex (mPFC), ventral hippocampus (vHPC) and cerebellum (see Figure 6A) and frozen at -80° until further use.

### 2.11 Quantitative Polymerase Chain Reaction (qPCR)

Gene expression was analysed using standardised qPCR methods with quantification using the 2^-ΔΔCt^ method^45^. Brain tissue from the mPFC, vHPC and the cerebellum was analysed. RNA was extracted using the RNeasy kit (QIAGEN) and was subsequently treated with DNAse to remove genomic DNA (TURBO DNA-free kit, Thermo Fisher Scientific). RNA was then converted to cDNA (RNA to cDNA EcoDry Premix, Random Hexamers, Clontech, Takara). cDNA samples were run in triplicate in 96 well reaction plates using SYBR-Green-based qPCR (SensiFast, HI-ROX, Bioline) according to manufacturer’s instructions using a StepOnePlus System (Applied Biosystems, Thermo Fisher Scientific). Genotypes were counterbalanced across plates and genes of interest were run alongside housekeeping genes *Gapdh* and *Hrpt1* for each sample, within the same reaction plate. All samples were run in triplicate and samples differing by >0.3 Cts were excluded. The change in expression of genes of interest, after normalisation to the two house-keeping genes (ΔCt) was transformed to yield 2^-ΔΔCt^ values. Relative changes from wildtype animals were calculated for each gene of interest.

### 2.12 Primers

Primers were designed to span at least one exon-exon junction and to match the target sequence only in mouse (Primer-Blast, NCBI) and were synthesised commercially (Sigma Aldrich). Primer efficiency was determined separately through a dilution series of cDNA samples from wildtype hippocampus, cerebellum and cortex. Primers with an efficiency between 90-110% were selected.

**Table 1.**
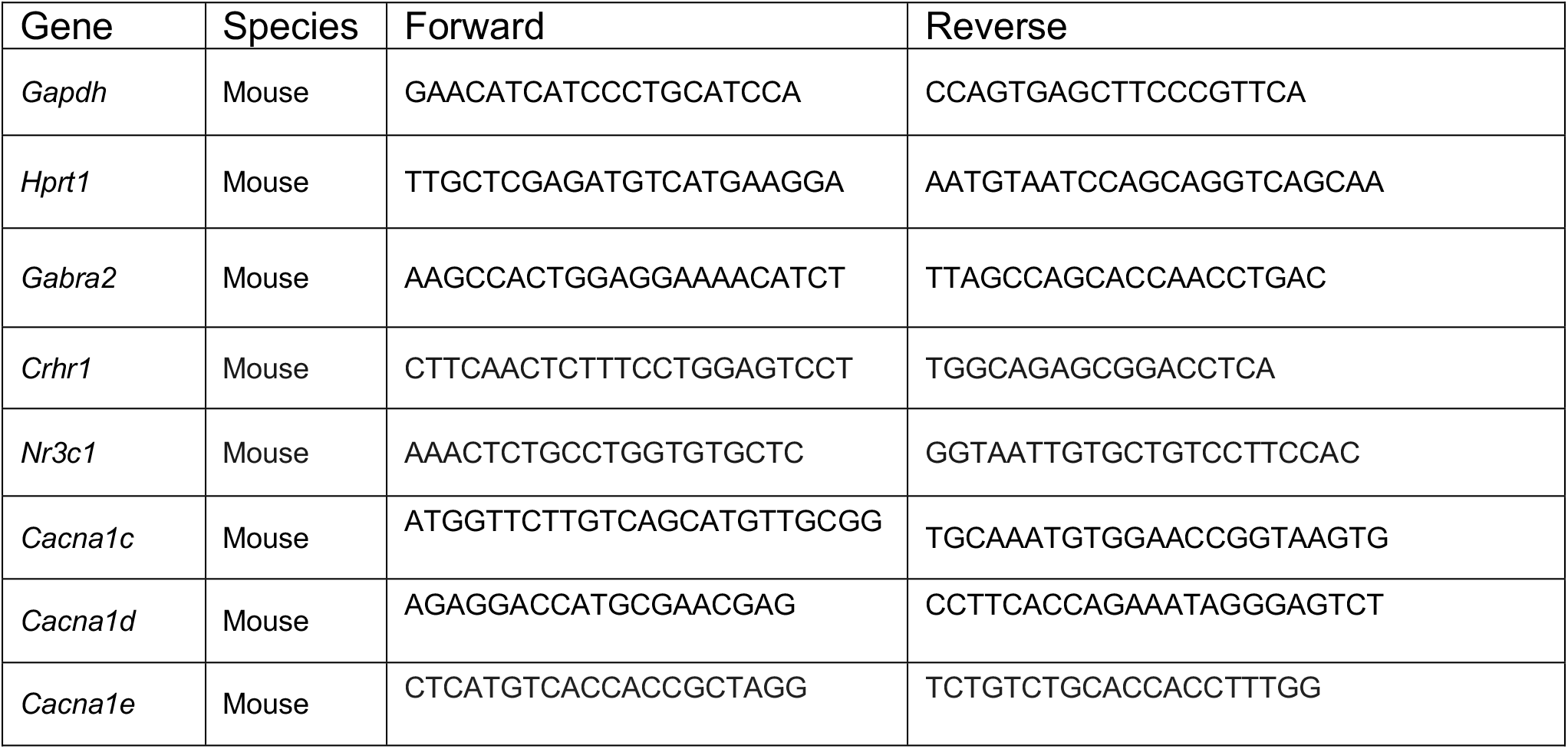
List of primer sequences used.

### 2.13 Genotyping

Genotyping was performed on post-mortem tail tip samples. Qiagen DNeasy Blood and Tissue Kits (Qiagen, Manchester, UK) were used to extract genomic DNA (gDNA) as per the manufacturers standard protocol. For *C3*^-/-^ mice, JAX protocol 27746 was used (common; ATCTTGAGTGCACCAAGCC, wildtype; GGTTGCAGCAGTCTATGAAGG, mutant; GCCAGAGGCCACTTGTATAG) and for *C3aR*^*-*/-^ JAX protocol 27638 was used (common; AGCCATTCTAGGGGCGTATT, wild type reverse; CATGGTTTGGGGTTATTTCG, mutant reverse; TTGATGTGGAATGTGTGCGAG). For both genotypes, a touchdown cycling protocol was used (see JAX protocols for details). Genotyping for *C5*^-/-^ mice was performed as described in ^38^.

### 2.14 Statistical analysis

All statistical analyses were carried out using GraphPad Prism 8.4.1 (GraphPad Software, CA, USA). Data was assessed for equality of variances using the Brown-Forsythe test and then appropriate parametric (*t* test, one-way or two-way ANOVA) or non-parametric (Kruskal-Wallis) tests used. *Post hoc* pairwise comparisons were performed using the Tukey or Dunn’s tests for parametric or non-parametric analyses, respectively. For all analyses, alpha was set to 0.05 and exact p values were reported unless p<0.0001. All p values were multiplicity adjusted^46^. Data are expressed as mean ± standard error of the mean.

The main between-subjects’ factor for all ANOVA analyses was GENOTYPE (WT, *C3*^-/-^, *C3aR*^-/-^, or *C5*^-/-^). For the EPM, LMA and FPS experiments, there were within-subject factors of ZONE (open, closed, middle), DAY (1,2,3) and STIMULUS INTENSITY (90, 100, 110 dB) respectively. Analysis of plasma corticosterone by two-way ANOVA included an additional between subject factor of CONDITION (baseline, EPM), and for the diazepam experiment, there was an additional between subject factor of DRUG (diazepam, vehicle). For qPCR analyses, ΔCt values were analysed by one-way ANOVA.

## 3. Results

### 3.1 Increased innate anxiety in *C3aR*^-/-^ but not *C3* ^-/-^ mice

Using a cohort of male wildtype, *C3*^-/-^ and *C3aR*^-/-^ mice we first assessed emotional reactivity in the elevated plus maze (EPM), a well-established test of innate anxiety in rodents which exploits the conflict between the drive to explore novel environments and the innate aversion towards open, brightly lit spaces^47,48^. Heatmaps indexing overall maze exploration over a 5-minute session demonstrated major differences in open arm exploration between genotypes (Figure 1A; see Supplementary Video 1 for representative examples). Notably, in comparison to wildtype and *C3*^*-*/-^ mice, *C3aR*^-/-^ mice took significantly longer to first enter the open arms (Figure 1B) and spent less time on the open arm per entry (Figure S2A), leading to a reduced overall duration spent in the open arms (Figure 1C), findings consistent with increased anxiety. The ethological parameters head dips and stretch attend postures (SAPs) also differed between genotypes (Figure 1D,E), with *C3aR*^-/-^ mice exhibiting decreases in the former and increases in the latter, a pattern of results again consistent with heightened anxiety^49^. We also noted a significantly increased frequency of head dipping in *C3*^*-*/-^ mice (Figure 1D), suggestive of reduced levels of anxiety relative to both wildtype and *C3aR*^-/-^ mice^50^.

**Figure 1.**
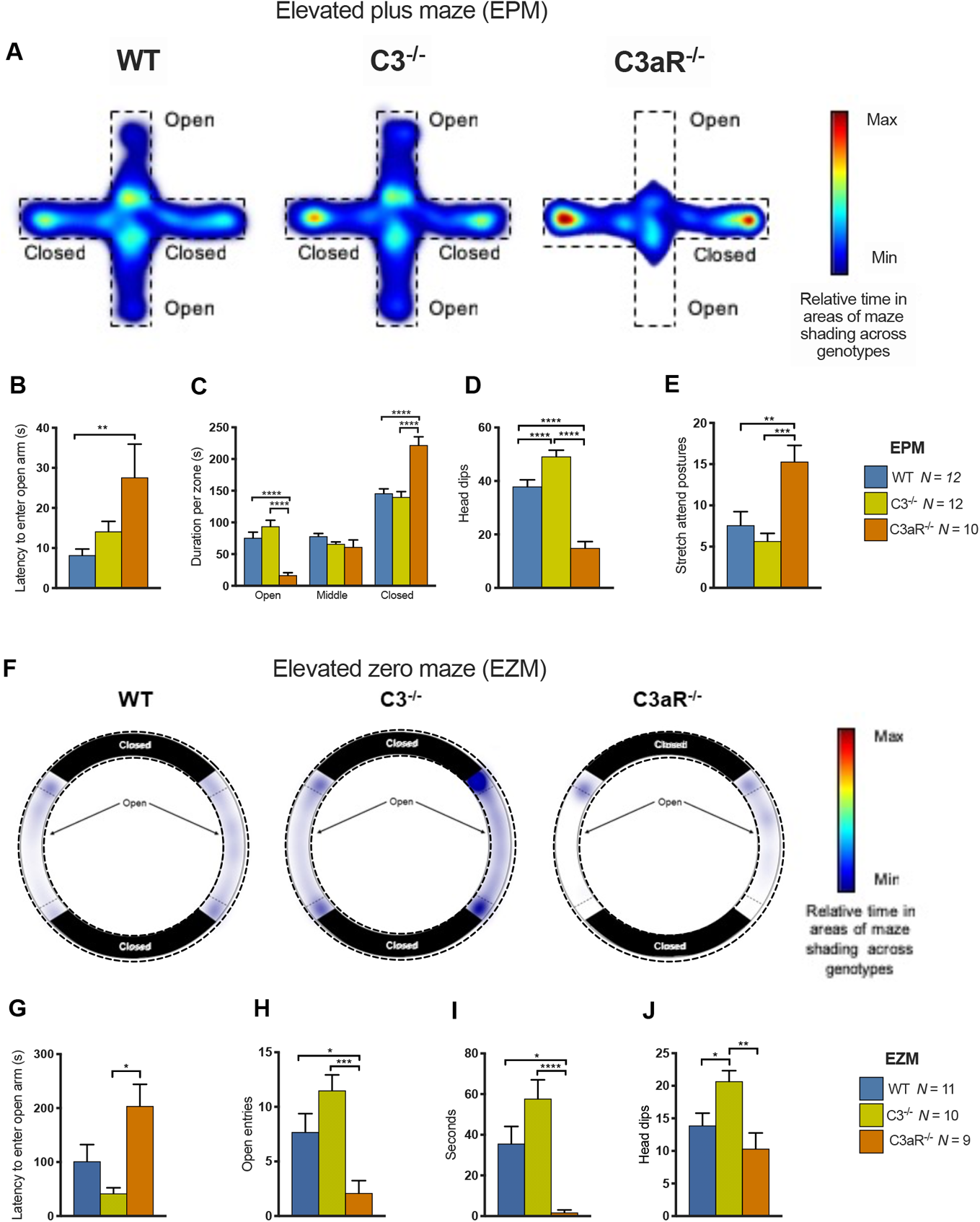
*C3aR*^-/-^, but not *C3*^-/-^ mice show increased anxiety-like behaviour in the elevated plus maze (EPM;A-E) and elevated zero maze (EZM;F-J). **(A)** Heatmaps displaying relative time per zone of the EPM across genotypes **(B)** Latency to first open arm visit; wildtype 8.21±1.53s, *C3*^*-*/-^ 14.1±2.52s, *C3aR*^-/-^ 27.6±8.31s, (H_2_=10.5, p=0.005). *Post hoc* tests demonstrated that *C3aR*^-/-^ mice took significantly longer to first enter the open arms than wildtype mice (p=0.0045). **(C)** *C3aR*^-/-^ mice distributed their time across the EPM differently to wildtype and *C3*^-/-^ mice (GENOTYPE×ZONE, F_4,62_=17.7, p=0.0001) spending less time in the open arms (*C3aR*^-/-^ 16.70±3.73s vs. wildtype 75.78±8.86s, p<0.0001, *C3aR*^-/-^ vs. *C3*^-/-^ 93.86±9.59s p<0.0001) and significantly more time in the closed arms (*C3aR*^-/-^ 221.88±12.06s vs. wildtype 146.01±7.01s, p<0.0001, and *C3*^-/-^ 140.04±8.61s p<0.0001). **(D)** *C3aR*^-/-^ (14.90±2.22) mice performed significantly fewer head dips than wildtype (37.92±2.53, p<0.0001) and *C3*^-/-^ mice (49.17±2.37, p<0.0001), whereas *C3*^-/-^ mice performed significantly more head dips than wildtype mice (p=0.0061; overall ANOVA F_2,31_=48.0, p<0.0001). **(E)** *C3aR*^-/-^ mice performed significantly more stretch attend postures (SAPs; 15.30±1.80) than wildtype (7.58±1.66, p=0.0042) and *C3*^-/-^ mice (5.67±0.94, p=0.0004; overall ANOVA F_2,31_=10.3, p=0.0004). **(F)** Heatmaps displaying relative exploration of the open segments of the elevated zero maze, across genotypes. Note that due to the height of the walls in the closed regions it was not possible to track mice or observe ethological behaviours such as grooming or SAPs. **(G)** There was a significant difference in the latency to first enter the open arms (wildtype 101.00±31.00s, *C3*^*-*/-^ 42.00±2.52s, *C3aR*^-/-^ 204.00±40.40s, H_2_=8.13, p=0.0171). *Post hoc* tests revealed that *C3aR*^-/-^ mice took significantly longer than *C3*^-/-^ mice to initially enter the open region (p=0.0140). **(H)** The number of entries made to open regions differed between genotypes (wildtype 7.69±1.69, *C3*^*-*/-^ 11.5±1.43s, *C3aR*^-/-^ 2.10±1.15, F_2,30_=8.96, p=0.0009). *C3aR*^-/-^ mice made significantly fewer entries to the open areas than wildtype (p=0.0324) and *C3*^-/-^ mice (p=0.0006) and **(I)** spent significantly less time on the open arms (1.77±1.29) compared to wildtype (35.7±8.43s, p=0.0132) and *C3*^*-*/-^ (57.7±9.32s, p<0.0001; overall Kruskal-Wallis test H_2_=19.2, p<0.0001). **(J)** *C3*^-/-^ mice performed significantly more head dips (20.7±1.62) than wildtype (13.9±1.89, p=0.048) and *C3aR*^-/-^ mice (10.3±2.42, p=0.0034; overall ANOVA F_2,27_=6.86,p=0.0039). Data are mean ± S.E.M. *, **, *** and **** represent p≤0.05, p≤0.01, p≤0.001 and p≤0.0001 for *post-hoc* genotype comparisons, respectively.

These initial data were consistent with an anxiogenic phenotype present in *C3aR*^-/-^ mice but absent in *C3*^-/-^ mice. We confirmed the findings in two further independent tests of anxiety using additional cohorts of animals. First we used the elevated zero maze (EZM, see Methods Section 2.6), another test of anxiety-like behaviour which similarly probes behavioural responses to exposed, illuminated spaces^51^. The data recapitulated the pattern of findings seen in the EPM (Figure 1F-J). Additional data from the open field test, where *C3aR*^-/-^ mice were more likely to avoid the centre of the arena, were also consistent in demonstrating a specific anxiety-like phenotype in *C3aR*^-/-^ but not *C3*^-/-^ mice (Figure S3). Given that several of the measures indexing anxiety were dependent on movement around the apparatus it was important to eliminate potential locomotor confounds. To address this issue, we measured activity independently in a non-anxiety provoking environment and found no differences in locomotor activity between genotypes (Figure S2C), demonstrating that anxiety measures were unlikely to be influenced by movement confounds. Importantly, experiments conducted in female mice demonstrated comparable *C3aR*^-/-^ anxiety phenotypes in both the elevated plus maze and open field (Figure S6&7).

### 3.2 Neuroendocrine response in *C3aR*^*-*/-^ and *C3*^-/-^ mice following exposure to the elevated plus maze

We next tested whether the behavioural measures of anxiety were paralleled by changes in plasma levels of the stress hormone corticosterone. In a separate cohort of wildtype, *C3*^-/-^ and *C3aR*^*-*/-^ male mice, we assayed plasma corticosterone 30 minutes after exposure to the EPM and compared levels to those of a group of animals who remained in their home-cages. There were no genotype differences in basal corticosterone levels; however, being placed on the EPM increased plasma corticosterone 6-15-fold in all genotypes, demonstrating that the EPM was a potent stressor (Figure 2A). *Post hoc* analyses showed a significantly greater EPM-evoked corticosterone response in the *C3aR*^-/-^ animals, consistent with their increased anxiety-like behaviour observed on the maze.

**Figure 2.**
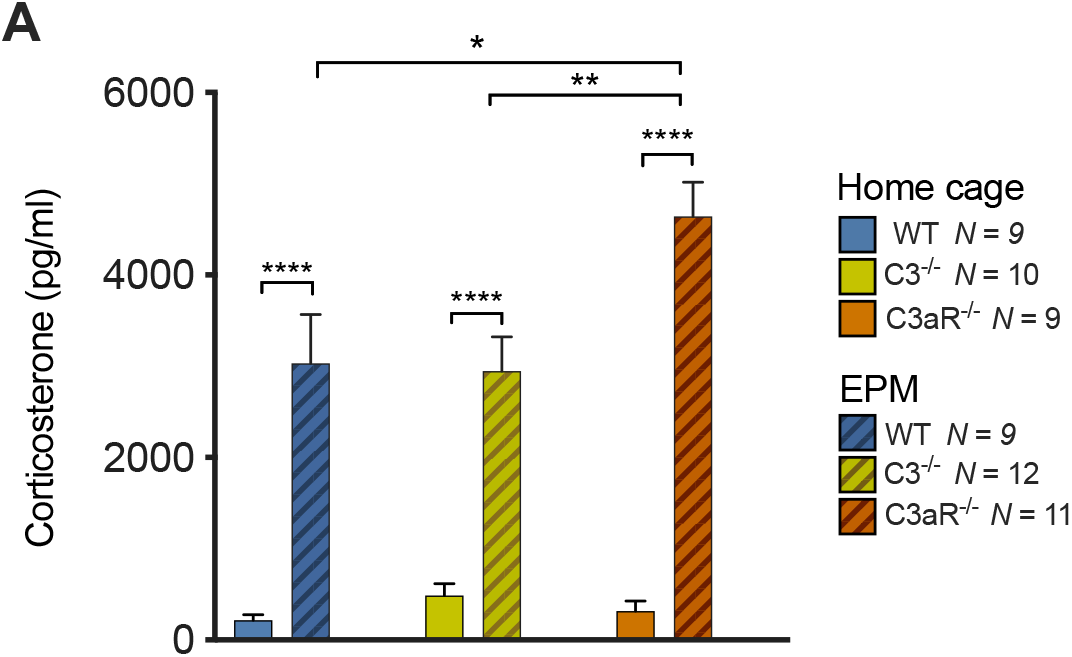
Neuroendocrine response following exposure to the elevated plus maze. **(A)** 5-minute exposure to the EPM significantly elevated corticosterone in all genotypes (main effect of CONDITION, F_1,54_=143, p<0.0001; baseline 344.66±63.70 vs. EPM 3553.84±274.13). There was a significant GENOTYPE × CONDITION interaction (F_2,54_=4.64, p=0.0138). *Post hoc* analysis showed that after the EPM, *C3aR*^-/-^ mice demonstrated significantly higher corticosterone levels (4640.27±376.13) than wildtype (3033.78±535.06, p=0.0127) and *C3*^-/-^ mice (2948.00±374.87, p=0.0032). *Post hoc* tests also indicated that there were no baseline differences between genotypes (wildtype 216.54±63.2 vs. *C3aR*^-/-^ 316.17±111.60 p>0.9999, wildtype vs. *C3*^-/-^ p=0.9927, and *C3*^-/-^ 485.60±130.35 vs.*C3aR*^-/-^ mice p=0.9992). Data represent mean + S.E.M. *, **, and **** represent p≤0.05, p≤0.01 and p≤0.0001 for *post-hoc* genotype comparisons, respectively.

### 3.3 Altered sensitivity of *C3aR*^-/-^ and *C3*^-/-^ mice to diazepam in the elevated plus maze

In a further independent cohort of male mice, we tested the sensitivity of EPM induced anxiety-like behaviour to the benzodiazepine diazepam, an established clinically effective anxiolytic drug^47,52^. Our initial behavioural findings were replicated in vehicle-treated animals across all behavioural indices of anxiety, again showing an anxiogenic phenotype in *C3aR*^-/-^ but not *C3*^-/-^ mice (Figure 3A). As anticipated, in wildtype mice 2mg/kg diazepam led to a trend for increased time on the open arms (Figure 3B) and a significant reduction in SAPs which are considered to reflect risk assessment behaviour^53-55^(Figure 3C). These effects were therefore consistent with reduced anxiety^50,55^. In contrast, the same dose of drug that was effective in eliciting anxiolysis in wildtypes was without effects in *C3aR*^-/-^ mice (Figure 3A,B,C) and produced a seemingly anxiogenic (increased anxiety) pattern of effects in *C3*^-/-^ mice (Figure 3A,B and Figure S4B,C). Locomotor activity monitored across all the maze (Figure S4D) indicated that wildtype and *C3aR*^-/-^ mice were unlikely to have been influenced by diazepam-induced sedation. In *C3*^-/-^ mice however, activity was significantly suppressed under drug conditions indicating a possible sedative effect. Together, these data indicated a fundamentally altered reactivity to diazepam in both *C3*^-/-^ and *C3aR*^-/-^ models.

**Figure 3.**
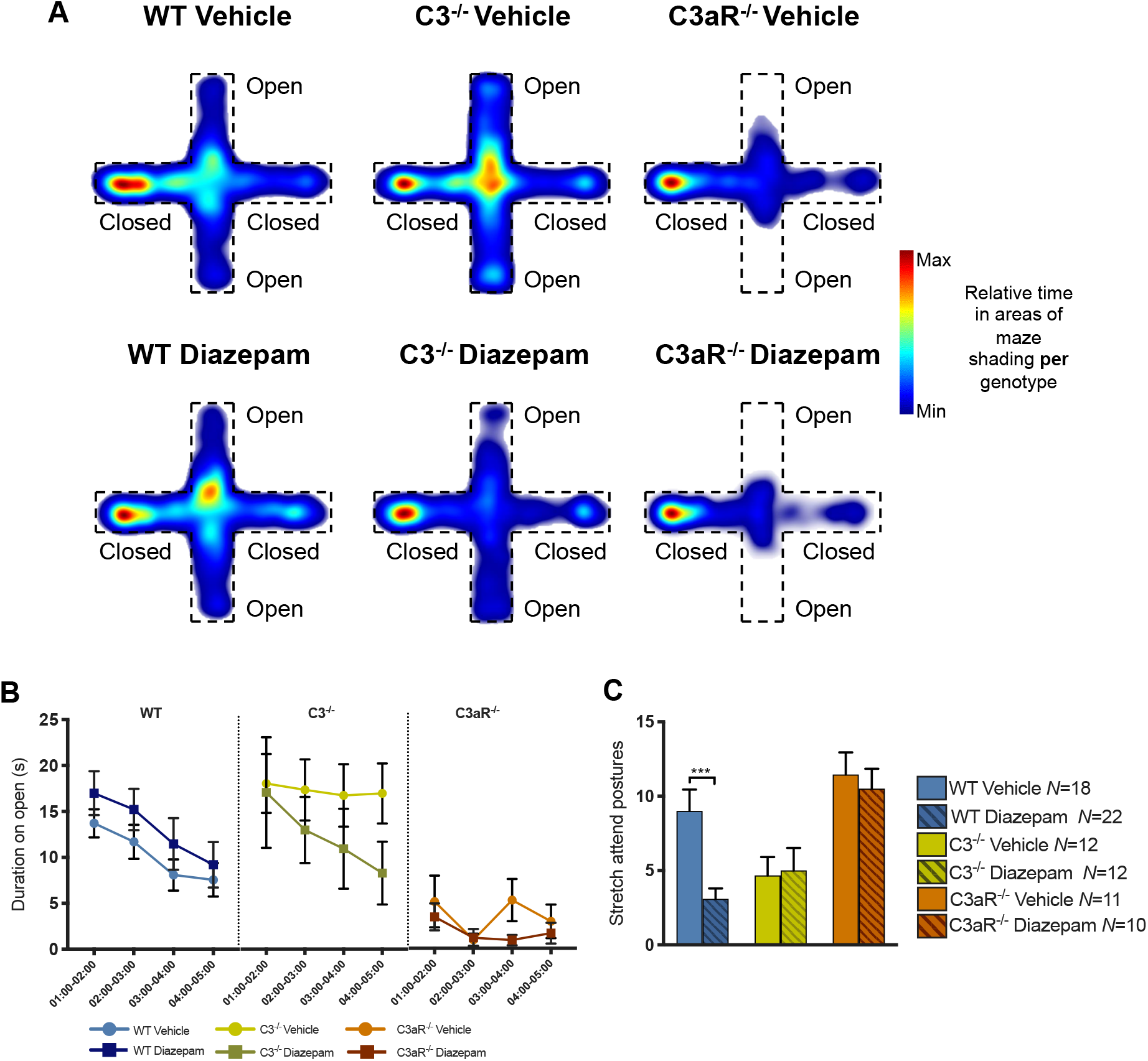
Altered sensitivity to diazepam in *C3aR*^-/-^ and *C3*^-/-^ mice. Behaviourally naïve mice were treated with either diazepam (2mg/kg, i.p) or vehicle injections once daily for 2 days and then 30 minutes prior to testing. (**A)** Heatmaps demonstrating duration spent in zones of the maze by vehicle treated and diazepam treated animals **(B)** Plots showing duration spent on open arms in 1-minute time bins (start-01:00 was excluded due to effect of diazepam in delaying initial entry to open arms across genotypes, see Supplementary Figure 4A). There was a trend for wildtype diazepam treated animals to spend more time on the open arms throughout the task although this did not reach significance (main effect of DRUG, F_1,38_=1.41, p=0.2462). In *C3*^-/-^ mice there was a strong tendency for drug treated animals to explore the open arms less than vehicle treated *C3*^-/-^ mice (main effect of DRUG, F_1,22_=1.25, p=0.2764). A similar, though less pronounced pattern was seen in *C3aR*^-/-^ mice (main effect of DRUG, F_1,19_=1.55, p=0.2284) **(C)** There were genotype differences in SAPs (main effect of GENOTYPE, F_2,79_=10.7, p<0.0001), a main effect of DRUG (F_1,79_=4.13, p=0.0454) and a significant GENOTYPE × DRUG interaction (F_2,79_=4.64, p=0.0138). *Post hoc* tests showed that diazepam significantly reduced the number of SAPs in wildtype mice only (wildtype vehicle 9.00±1.44 vs. wildtype diazepam 3.09±0.71, p=0.0006, *C3*^-/-^ vehicle 4.67±1.24 vs. *C3*^-/-^ diazepam 5.00±1.51, p=0.9975, *C3aR*^-/-^ vehicle 11.45±1.49 vs. *C3aR*^-/-^ diazepam 10.50±1.34, p=0.9558). Data are mean + S.E.M. *, **, and *** represent p≤0.05, p≤0.01 and p≤0.001 for *post-hoc* genotype comparisons, respectively.

### 3.4 Enhanced fear learning in *C3*^-/-^ but not *C3aR*^-/-^ mice

Psychiatric disorders are associated with maladaptive responses to both innate and learned aversive stimuli^56,57^. We therefore extended our analysis to investigate whether the behavioural dissociations in innate anxiety observed between *C3*^-/-^ and *C3aR*^-/-^ mice would also apply to learned or conditioned fear, where a previously neutral cue generates a fear response as a result of predicting an aversive outcome. In a further group of male mice, we used the fear-potentiated startle (FPS) paradigm^30,58^ a well-established method of generating learned fear responses to an acute and imminent danger signal that is characteristic of fear. In this paradigm (see Figure 4A and Methods Section 2.7) fear learning is indexed by an enhanced response to a startling noise in the presence of a cue (the conditioned stimulus or CS) previously paired with mild foot shock (the unconditioned stimulus). In the pre-conditioning session, pulse-alone trials revealed increased basal startle reactivity in both *C3aR*^-/-^ and *C3*^-/-^ mice relative to wildtype (Figure 4B). Increased reactivity to the unconditioned foot shock stimulus (in the absence of any startle stimulus) during the conditioning session was also observed in both knockouts (Figure 4C). However, these common effects of genotype were not seen in the fear-potentiated startle measures which index fear learning. Whilst all groups showed the expected enhancement of the startle response in the presence of the CS, the effect of the CS was significantly greater in *C3*^-/-^ animals relative to the *C3aR*^-/-^ and wildtype mice (Figure 4D), indicating enhanced learning of the fear related-cue by the *C3*^-/-^ mice. This pattern of effects was also observed in female mice (Figure S8). This was the opposite pattern of effects to those observed in the tests of innate anxiety and showed a double dissociation in the impact of manipulating C3 and C3aR function that depended fundamentally on the nature of the aversive stimulus.

**Figure 4.**
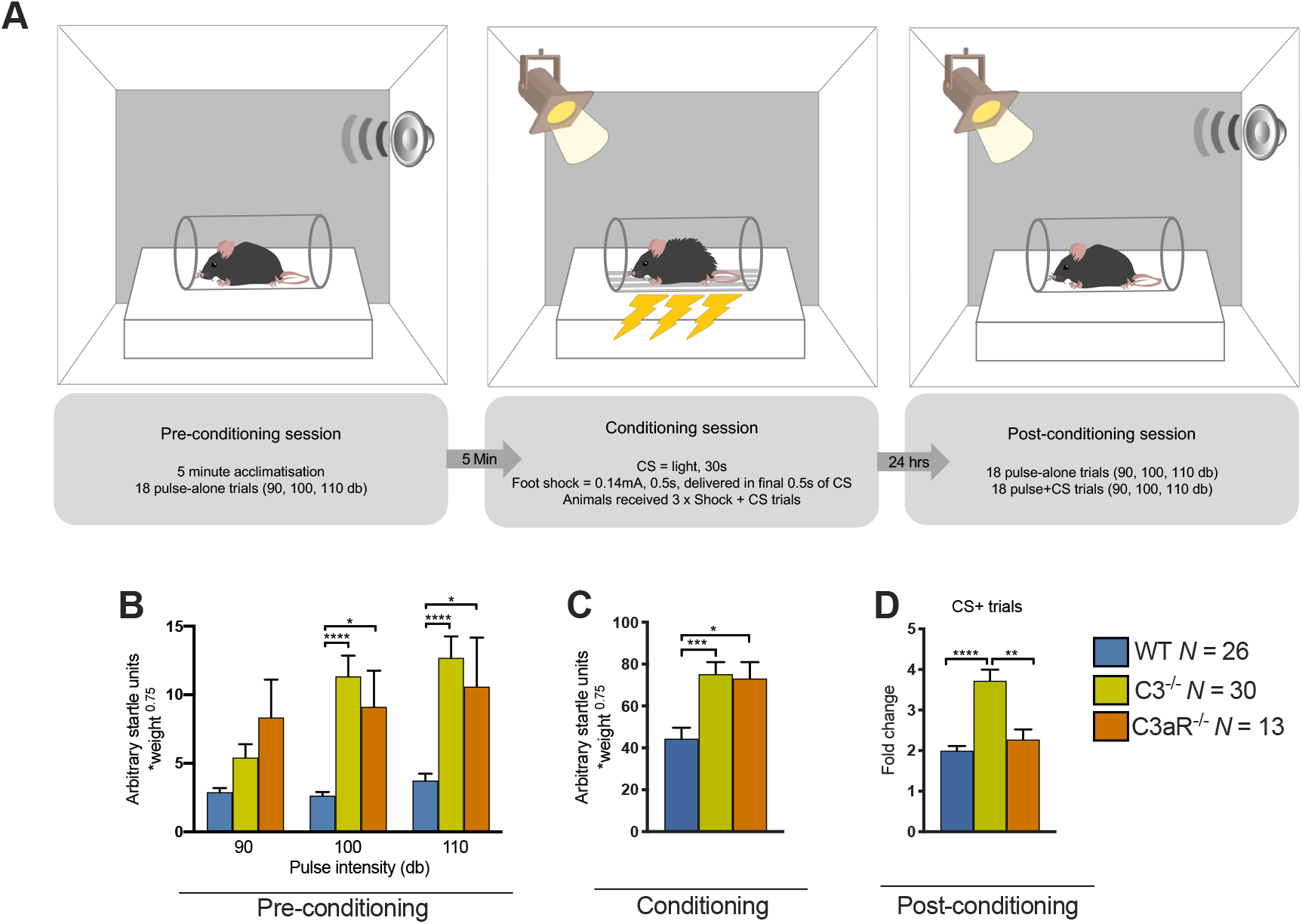
Enhanced fear-potentiated startle in *C3*^-/-^ but not *C3aR*^-/-^ mice. **(A)** Flow chart depicting the FPS protocol used, which took place in three separate sessions over two consecutive days. Baseline startle reactivity to a range of pulse intensities was assessed in the pre-conditioning session, immediately preceding the conditioning session in which a visual stimulus (light) was paired with 3 weak foot shocks. 24 hours later, subjects were re-introduced to the same chamber and startle reactivity was compared between pulse-alone trials and pulse+CS trials to determine the degree of FPS. On all trials, the peak startle response was recorded and normalised for body weight differences using Kleiber’s 0.75 mass exponent, and fold-changes calculated. **(B)** There was a significant main effect of GENOTYPE (F_2,66_=9.04, p=0.0003) and a significant GENOTYPE × STIMULUS INTENSITY interaction (F_4,132_=7.55, p<0.0001). *C3*^-/-^ and *C3aR*^*-/-*^ mice demonstrated increased levels of startle responding relative to wildtype mice at 100dB (*C3*^-/-^ 11.34±1.51 vs. wildtype 2.63±0.26, p<0.0001, *C3aR*^*-/-*^ 9.12±2.63 vs. wildtype p=0.0174) and 110dB (*C3*^-/-^ 12.69±1.55 vs. wildtype 3.74±0.50, p<0.0001, *C3aR*^*-/-*^ 10.58±3.58 vs. wildtype p=0.0111) **(C)** *C3*^-/-^ and *C3aR*^-/-^ mice also showed increased startle responding to the footshock+CS (*C3*^-/-^ 75.18±5.73, *C3aR*^-/-^ 73.14±7.78) pairings relative to wildtype mice (44.34±5.29, *C3*^-/-^ vs. wildtype p=0.0006, *C3aR*^-/-^ vs. wildtype p=0.0137, overall ANOVA F_2,66_=8.7, p=0.0004), although it should be noted that responses were much greater to these stimuli in all mice than to the startle stimuli in the pre-conditioning session. **(D)** In the post-conditioning session, all mice demonstrated increases to the pulse+CS stimuli in comparison to pulse-alone stimuli, as demonstrated by the fold-change increase in startle responding, however, this effect was significantly increased in *C3*^-/-^ mice (3.72±0.27) relative to wildtype (1.99±0.11, p<0.0001) and *C3aR*^-/-^ mice (2.27±0.24, p=0.0056, overall Kruskal-Wallis test H_2_=27.7, p<0.0001). Data are mean + S.E.M. *, **, *** and **** represent p≤0.05, p≤0.01, p≤0.001 and p≤0.0001 for *post-hoc* genotype comparisons, respectively.

### 3.5 Complement signalling pathways underlying abnormal learned fear phenotypes in *C3*^-/-^ mice

Given the central role of C3 within the complement system, its deletion affects a number of distal pathways (Figure 5A), with the activity of the C3a/C3aR, C3b/CR3, C5a/C5aR and terminal pathways affected. Therefore, the loss of function of any of these pathways may have contributed to the observed fear learning phenotype in *C3*^-/-^ mice. However, it is possible to exclude effects due to loss of the C3a/C3aR pathway since the fear learning phenotype was specific to *C3*^-/-^ and not *C3aR*^-/-^ mice (Figure 4D). This left iC3b/CR3 signalling and/or pathways downstream of C5 (i.e. C5a/C5aR, terminal pathway) as the remaining possibilities. In order to distinguish between these pathways, we repeated the FPS experiment with the addition of *C5*^-/-^ mice. This model has intact C3a/C3aR and iC3b/CR3 signalling, but lacks C5a/C5b and terminal pathway activity, as do *C3*^-/-^ mice (Figure 5A). We hypothesised that if C*5*^-/-^ mice also displayed enhanced fear-potentiated startle, then the phenotype in *C3*^-/-^ mice would likely be due to a loss of C5a/C5aR signalling or the terminal pathway. On the other hand, if *C5*^-/-^ mice demonstrated normal fear-potentiated startle, this would confine the likely mediating pathway in *C3*^-/-^ mice to iC3b/CR3 signaling (Figure 5A).

**Figure 5.**
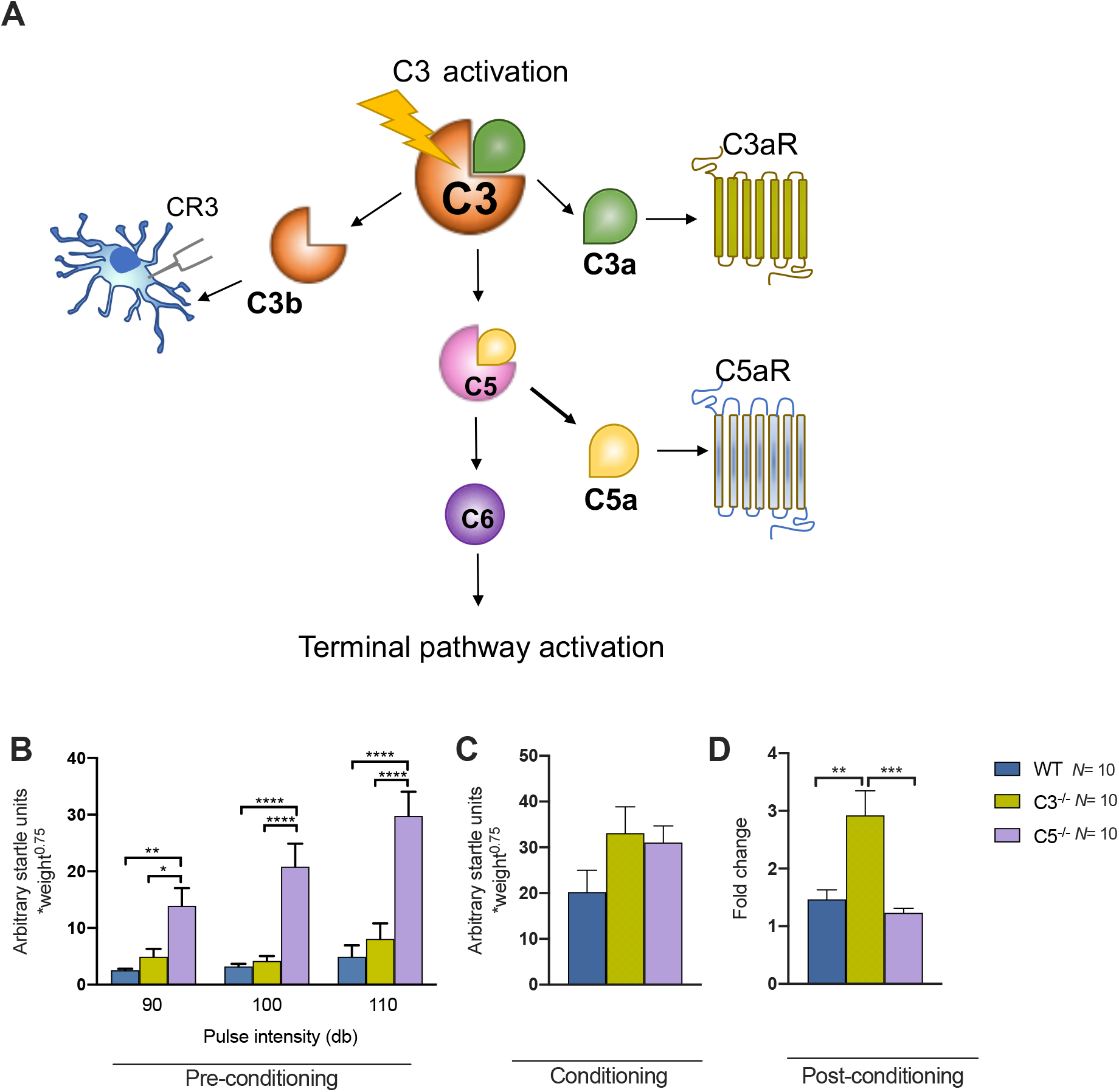
Pathways underlying fear learning phenotypes in *C3*^-/-^ mice. **(A)** C3 activation leads to generation of cleavage fragments C3a and C3b. The former signals via C3aR whereas the latter signals via complement receptor 3 (CR3). C3b is also necessary for forming the convertase enzyme that cleaves C5. Upon cleavage, C5 generates the fragments C5a and C5b (not shown). C5a signals via the C5aR, whereas C5b propagates activity of the terminal complement pathway via C6. Since C3 cannot be activated in *C3*^-/-^ mice, the action of all these pathways (C3a/C3aR, C3b/CR3, C5a/C5aR, terminal pathway) is absent. By using *C5*^-/-^ mice, which lack C5a/C5aR and terminal pathway activity, we examined whether lack of C3b/CR3, C5a/C5aR or the terminal pathway was responsible for fear learning phenotypes in *C3*^-/-^ mice. **(B)** In the pre-conditioning session there were significant main effects of GENOTYPE (F_2,27_=18.4, p<0.0001) and STIMULUS INTENSITY (F_2,54_=19.0, p<0.0001) and a significant GENOTYPE × STIMULUS INTENSITY interaction (F_4,54_=7.00, p<0.0001). *C5*^-/-^ mice demonstrated increased levels of startle responding relative to wildtype and *C3*^-/-^ mice at all stimulus intensities (90dB; WT 2.55±0.26 vs. *C5*^-/-^ 13.92±3.14, p=0.0069, *C3*^-/-^ 4.92±1.40 vs. *C5*^-/-^ p=0.0405, WT vs. *C3*^-/-^ p=0.7919; 100dB; WT 3.23±0.45 vs. *C5*^-/-^ 20.83±4.07, p<0.0001, *C3*^-/-^ 4.924.17±0.88 vs. *C5*^-/-^ p<0.0001, WT vs. *C3*^-/-^ p=0.9639; 110dB; WT 4.92±2.03 vs. *C5*^-/-^ 29.78±4.29, p<0.0001, *C3*^-/-^ 8.07±2.76 vs. *C5*^-/-^ p<0.0001, WT vs. *C3*^-/-^ p=0.6643). **(C)** There were no significant differences in startle responses to the footshock+CS pairings during the conditioning session (WT 20.23±4.76, *C3*^-/-^ 33.10±5.74, *C5*^-/-^ 31.08±3.59, F_2,27_=2.10, p=0.1421). **(D)** In the post-conditioning session, all mice demonstrated increases to the pulse+CS stimuli in comparison to pulse-alone stimuli, as demonstrated by the fold-change increase in startle responding, however, this effect was again significantly increased only in *C3*^-/-^ mice (2.92±0.43) relative to wildtype (1.47±0.17, p=0.0020) and *C5*^-/-^ mice (1.23±0.08, p=0.0004, overall ANOVA F_2,27_=11.5, p=0.0002). Data represent mean + S.E.M. *, **, *** and **** represent p≤0.05, p≤0.01, p≤0.001 and p≤0.0001 for *post-hoc* genotype comparisons, respectively.

Results from the pre-conditioning session demonstrated increases in the startle response of C*5*^-/-^ mice, (Figure 5B) although in this instance the previously observed enhanced startle reactivity in C*3*^-/-^ mice (Figure 4B) was not replicated. In the conditioning session, there was again evidence of increased startle responses to shock (Figure 5C) in C*3*^-/-^ mice and responses were of a similar magnitude in C*5*^-/-^ mice, although these were not significantly different to wildtype. We replicated the previous finding of enhanced fear-potentiated startle in *C3*^-/-^ mice (Figure 5D), but critically both male (Figure 5) and female (Figure S9) *C5*^-/-^ mice showed no evidence of enhanced fear learning and were comparable to wildtypes, indicating that loss of iC3b/CR3 signalling, but not loss of C5a/C5aR and the terminal pathway, was involved in the *C3*^-/-^ fear learning phenotype. Additionally, we did not observe innate anxiety-like phenotypes in *C5*^-/-^ male mice (Figure S5).

This pattern of effects allowed us to distinguish between the likely mechanisms underlying the enhanced learned fear in *C3*^-/-^ mice, as we could exclude concomitant loss of C5a/C5aR signalling or molecules downstream of C5, and hence also exclude an explanation based on effects of C5a/C5aR signalling on developmental neurogenesis^59,60^. Instead, these data raised the possibility of an explanation based on the established effects of the iC3b/CR3 pathway on synaptic pruning, a mechanism involving microglia mediated elimination of synapses impacting on neurodevelopment and learning-related synaptic plasticity^13,14,61^.

### 3.6 Differential expression of stress and anxiety related genes in *C3aR*^*-*/-^ and *C3*^-/-^ mice

We next sought to assess whether the dissociations in innate anxiety and learned fear between *C3aR*^*-*/-^ and *C3*^-/-^ models were associated with differential gene expression in brain regions associated with emotional behaviours. We assayed gene expression in male mice, within three regions of recognised importance in stress and anxiety; the medial prefrontal cortex (mPFC), ventral hippocampus (vHPC) and cerebellum^28,62^(Figure 6A). Given our previous data showing differential corticosterone responses and altered sensitivity to diazepam in both knockouts, we first measured expression of the glucocorticoid receptor *Nr3c1* and the corticotrophin-releasing hormone receptor 1 *Crhr1*, together with *Gabra2* which encodes the GABA_A_ receptor α2 subunit responsible for mediating benzodiazepine anxiolysis^63^. There were no effects of genotype on *Crhr1* and *Gabra2* mRNA expression in any of the brain regions assayed (Figure 6B,C). *Nr3c1* expression did however show effects of genotype with trends indicating increases in *C3aR*^-/-^ mice in the mPFC and vHPC, and significantly increased expression in cerebellum that was common to both knockouts (Figure 6D).

**Figure 6.**
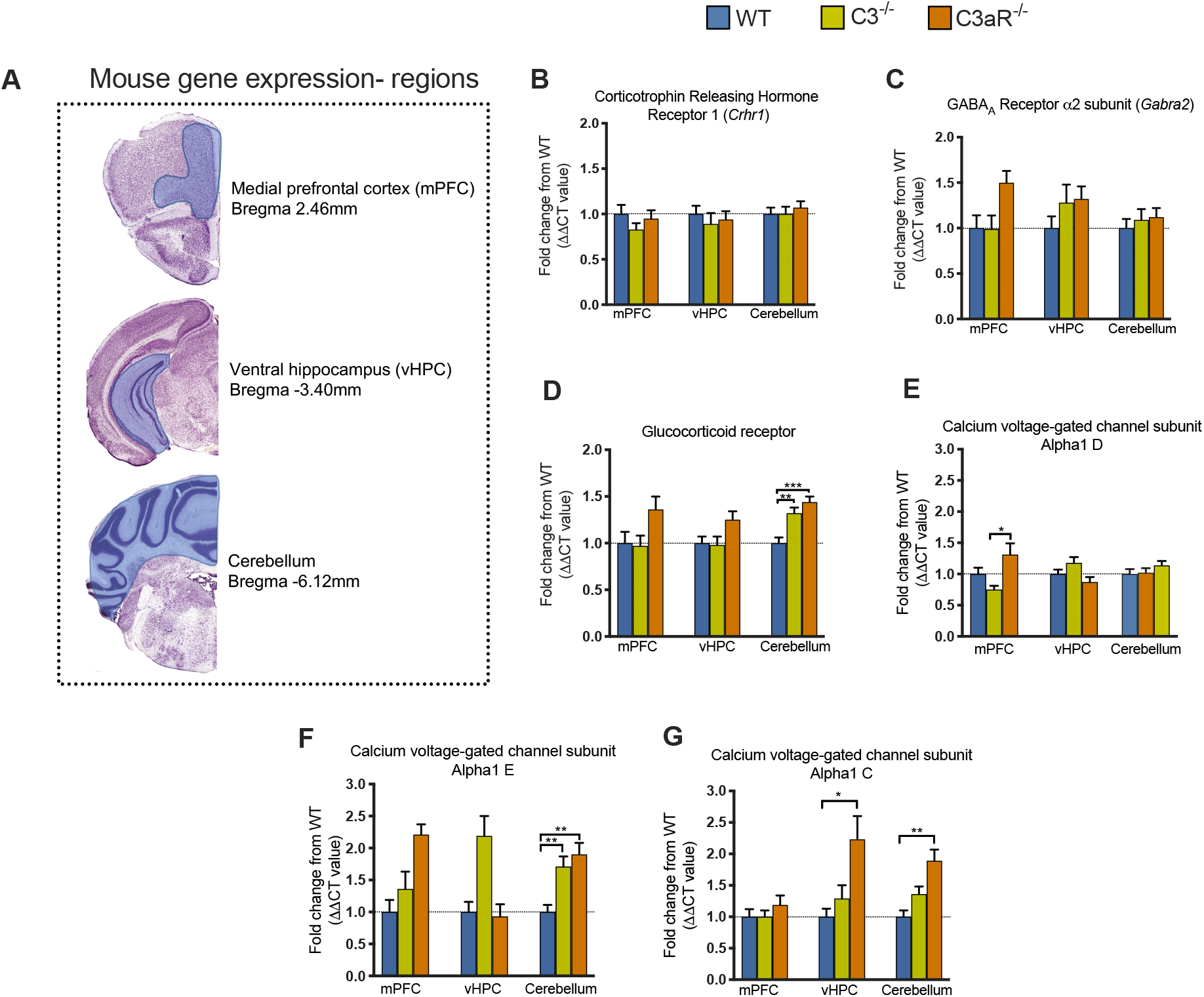
mRNA expression of stress and anxiety related genes. **(A)** Animals were culled and the medial prefrontal cortex (mPFC), ventral hippocampus (vHPC) and cerebellum were dissected. **(B)** There were no significant differences in the expression of Corticotrophin releasing hormone receptor 1 (*Crhr1*) in any region, across genotypes (mPFC F_2,53_=0.587, p=0.5597, N wildtype=20, *C3*^-/-^=17, *C3aR*^-/-^= 19; vHPC F_2,49_=0.169, p=0.8450, N WT=20, *C3*^-/-^=15, *C3aR*^-/-^= 17; cerebellum F_2,47_=0.0482, p=0.8346, N WT=19, *C3*^-/-^=17, *C3aR*^-/-^= 14). **(C)** There were also no significant changes in the expression of the GABA_A_ receptor α2 subunit (*Gabra2*) in any region, across genotypes (mPFC H_2_=1.04, p=0.5939, N wildtype=20, *C3*^-/-^=19, *C3aR*^-/-^= 16; vHPC F_2,49_=0.721, p=0.4914, N WT=20, *C3*^-/-^=13, *C3aR*^-/-^= 19; cerebellum F_2,47_=0.221, p=0.8026, N WT=18, *C3*^-/-^=17, *C3aR*^-/-^= 15). **(D)** Expression of the glucocorticoid receptor (*Nr3c1*) was significantly increased in the cerebellum of both *C3*^-/-^ and *C3aR*^-/-^ groups (F_2,61_=10.3, p=0.0002, *C3*^-/-^ vs. wildtype p=0.0023, *C3aR*^-/-^ vs. wildtype p=0.0002, N wildtype=19, *C3*^-/-^=20, *C3aR*^-/-^= 15). There were trends towards increased expression of the glucocorticoid receptor gene *Nr3c1* in the mPFC and vHPC of *C3aR*^-/-^ mice but these were not significant (mPFC; F_2,56_=1.33, p=0.2723, N wildtype=20, *C3*^-/-^=20, *C3aR*^-/-^= 19, vHPC; F_2,62_=1.11, p=0.3345, N wildtype=23, *C3*^-/-^ =20, *C3aR*^-/-^= 22). **(E)** Calcium voltage-gated channel subunit Alpha 1d (*Cacna1d*) expression was changed in the mPFC (F_2,36_=7.52, p=0.0407) owing to altered expression between *C3*^-/-^ and *C3aR*^-/-^ mice (p=0.0314; N wildtype=11, *C3*^-/-^=13, *C3aR*^- /-^= 15). There were no differences in the vHPC (F_2,31_=2.27, p=0.1199, N wildtype=14, *C3*^-/-^=10, *C3aR*^-/-^= 10) or cerebellum (F_1,39_=0.648, p=0.5286, N wildtype=14, *C3*^-/-^=16, *C3aR*^-/-^= 12). **(F)** Expression of the Calcium voltage-gated channel subunit Alpha 1e (*Cacna1e*) was significantly upregulated in the cerebellum of both knockouts (F_2,39_=7.52, p=0.0017, wildtype vs. *C3*^-/-^ p=0.0082, wildtype vs. *C3aR*^-/-^ p=0.0032; N wildtype=14, *C3*^-/-^=16, *C3aR*^-/-^= 12). There were borderline significant changes in expression in the vHPC (F_2,32_=3.15, p=0.0565, N wildtype=14, *C3*^-/-^=11, *C3aR*^-/-^= 10) and no significant changes in the mPFC (H_2_=3.43, p=0.1802, N wildtype=11, *C3*^-/-^=12, *C3aR*^-/-^= 15).**(G)** Expression levels of the Calcium voltage-gated channel subunit Alpha 1c (*Cacna1c*) were significantly increased in *C3aR*^-/-^ mice in a regionally specific manner in the vHPC (F_2,47_=3.20, p=0.0496, *C3*^-/-^ vs. wildtype p=0.6895, *C3aR*^-/-^ vs. wildtype p=0.0295, N wildtype=21, *C3*^-/-^=13, *C3aR*^-/-^= 16) and the cerebellum (F_2,54_=5.84, p=0.0051, *C3*^-/-^ vs. wildtype p=0.1613, *C3aR*^-/-^ vs. wildtype p=0.0024, N wildtype=20, *C3*^-/-^=20, *C3aR*^-/-^= 17). There were no significant changes in the mPFC (F_2,52_=1.04, p=0.5939, N wildtype=20, *C3*^-/-^=19, *C3aR*^-/-^= 16). Data represent fold change from wildtype + SEM. *, **, *** represent p≤0.05, p≤0.01 and p≤0.001 for *post-hoc* genotype comparisons, respectively.

As activation of C3aR has been shown to stimulate calcium influx from the extracellular space^2-4,64,65^ and calcium channel subunit variants have strong links to risk for psychiatric and neurological disorders, as well as anxiety phenotypes^66,67^, we also investigated a panel of voltage-gated calcium channels. Expression of *Cacna1d*, which encodes the Cav1.3 channel of L-type calcium gated voltage channels, was increased in the mPFC of *C3aR*^-/-^ mice (Figure 6E) and in both *C3*^-/-^ and *C3aR*^-/-^ mice there was upregulation of cerebellar *Cacna1e*, which encodes the Cav2.3 channel of R-type voltage gated calcium channels (Figure 6F). We also investigated the expression of *Cacna1c* which encodes the Ca_v_1.2 subunit of L-type voltage gated calcium channels that forms the channel pore allowing calcium entry^68^. We found genotype and brain region specific changes in *Cacna1c* expression, with selective increases in expression in *C3aR*^-/-^ mice in the vHPC and cerebellum, but not the mPFC (Figure 6G).

## 4. Discussion

We have used knockout models manipulating specific complement proteins to reveal dissociable effects of complement pathways on distinct elements of aversive behaviours. A main finding was a profound innate anxiety phenotype in the *C3aR*^-/-^ mice that was absent in *C3*^-/-^ mice. The specificity of the anxiety phenotype exhibited by *C3aR*^-/-^ mice at the behavioural level was paralleled by EPM-evoked corticosterone levels, confirming the validity of the EPM as an index of anxiety-like behaviour. A second main finding was a double dissociation in terms of fear learning, whereby *C3*^-/-^ and not *C3aR*^-/-^ mice displayed enhanced conditioned fear reactivity. Using *C5*^-/-^ mice, we conclude that the mechanisms accounting for the fear learning phenotype in *C3*^-/-^ mice are likely to arise from lack of iC3b/CR3 pathway activity. These data indicate distinct effects mediated by closely related elements of the complement system that differentially impact upon the neural mechanisms underlying innate anxiety and learned fear.

Our findings extend previous findings of abnormal anxiety behaviour in *C3aR*^-/-^ mice^35^. Our use of specific complement knockout models allowed us to further pinpoint the likely complement pathways and potential mechanisms underlying C3aR-mediated anxiety. Since C3a is solely produced via C3 cleavage, and C3aR is the canonical receptor for C3a, we expected that any phenotypes dependent on the binding of C3a to the C3aR would be apparent in both *C3*^-/-^ and *C3aR*^-/-^ models. However, this was not borne out in our data. Given the divergence in phenotypes seen, one explanation is that the *C3aR*^-/-^ anxiety phenotypes are independent of C3a and instead mediated by an alternative ligand. It has long been speculated that there may be promiscuity of the C3aR due to its unusually large second extracellular loop^69^. Indeed, a cleavage fragment of the neuropeptide precursor protein VGF (non-acronymic), TLQP-21, was recently reported to bind the C3aR^70,71^. This peptide has pleiotropic roles including in the stress response^72^ and its precursor VGF is widely expressed throughout the CNS^73^ and in regions associated with stress reactivity such as the hypothalamus, where there is evidence for C3aR expression^74,75^. Determining whether the mechanisms underlying innate anxiety phenotypes in *C3aR*^-/-^ mice are dependent on TLQP-21/C3aR interactions will be a priority for future work.

Whether the *C3aR*^-/-^ phenotypes described here are the result of ongoing effects of *C3aR* deletion in the adult brain or instead the enduring consequence of neurodevelopmental impacts of *C3aR* deficiency also remains to be determined. On the basis of previous findings implicating C3aR in both developmental neurogenesis^76^ and in acute brain changes following behavioural manipulations^77^, both are possibilities. One strategy would be to test whether acute administration of the C3aR antagonist SB290157^78^ phenocopies the constitutive knockout of *C3aR*, though at present no data is available on the CNS penetration of SB290157 and this molecule has received criticism due to evidence of agonist activity^79^.

We also probed mechanisms underpinning the *C3aR*^-/-^ innate anxiety phenotype by assessing the effects of the anxiolytic drug diazepam. We found that a dose of diazepam that was anxiolytic in wildtype mice had no effect in *C3aR*^-/-^ mice. Stretch attend postures, thought to reflect risk assessment behaviour^54^, are highly sensitive to pharmacological manipulation^50,80^ and in agreement with our own findings, diazepam has been shown to specifically decrease SAPs in the absence of effects on open arm exploration^81^. Importantly, *C3aR*^-/-^ mice consistently performed more SAPs than other genotypes, and therefore floor effects cannot account for the pattern of results observed. Benzodiazepines act on GABA_A_ receptors^63^ however we found no significant changes in expression of *Gabra2*, a GABA_A_ receptor subunit responsible for anxiolytic actions of benzodiazepines in tests of innate anxiety^63^, in the brain regions sampled. This raises the possibility of alternative molecular mechanisms mediating the anxiety phenotypes seen in the *C3aR*^-/-^ model. Whatever the molecular underpinnings of the dissociable anxiety phenotypes, our data show a profoundly altered effect of diazepam in both knockouts; a lack of response in *C3aR*^-/-^ and an apparent paradoxical anxiogenic effect of the drug in *C3*^-/-^ mice, though this interpretation needs to take into account an apparent selective sedative effect in *C3*^-/-^ mice.

In contrast to innate anxiety, we observed a specific effect of *C3* knockout on conditioned fear. The absence of a comparable phenotype in *C3aR*^-/-^ and *C5*^-/-^ mice suggested that these effects were unlikely to be due to loss of either C3a/C3aR, C5a/C5aR, or the terminal pathway, and instead that enhanced fear learning phenotypes in *C3*^-/-^ mice were likely dependent on loss of the iC3b/CR3 pathway. This pathway has been strongly implicated in activity dependent synaptic elimination during neurodevelopment^13,25^ and in age-dependent synapse loss^82^. While demonstrations of complement mediated synaptic pruning during development have centered on the visual system, complement-mediated microglial phagocytosis of dopamine D1 receptors has been demonstrated in the nucleus accumbens with functional impacts on social behaviour^83^. It remains to be seen whether complement mediated processes of this nature, within brain regions linked to fear processing such as the ventral hippocampus, amygdala and prefrontal cortex are responsible for enhanced fear learning in *C3*^-/-^ mice, or whether this phenotype is a general consequence of altered synaptic elimination throughout the *C3*^-/-^ brain. In addition to developmental processes, the iC3b/CR3 pathway could also be involved acutely in fear learning. *C3* mRNA is upregulated during discrete stages of fear learning^77^ and microglial CR3 is implicated in long term depression^84^. Furthermore, complement-mediated engulfment of synapses by microglia may be important in the forgetting of fear memories^14^.

At the gene expression level, we found some changes which were common to both knockouts, and one result that was specific to *C3aR*^-/-^ mice. Regarding the latter, there was a highly specific increase in expression of *Cacna1c* in the ventral hippocampus and cerebellum of *C3aR*^-/-^ mice. This finding is of potential interest given the strong evidence implicating *CACNA1C* variants in genetic risk for a broad spectrum of psychiatric disorders including schizophrenia and bipolar disorder^66^, with anxiety phenotypes reported in both human risk variant carriers^85^ and animal models^86-89^. Furthermore, recent evidence indicates convergent polygenic mechanisms shared between complement and other psychiatric risk genes^90^, including calcium regulation pathways, and thus our study lends further support to an interaction between these systems. Whether alterations in *Cacna1c* are of direct functional relevance to the *C3aR*^-/-^ anxiety phenotypes observed here remains to be determined experimentally. We also observed increased cerebellar expression of the glucocorticoid receptor in both *C3*^-/-^ and *C3aR*^-/-^ mice, suggesting that these alterations may result from the absence of C3a/C3aR signalling. Expression of these genes did not differentiate between models and therefore were unlikely to contribute to the dissociable effects of the knockouts on behaviour and stress hormone physiology. Future studies of neuronal activity in brain regions linked to emotion may be more informative in terms of functional neuroanatomy underlying the anxiety-related behavioural and physiological differences seen in the knockout models.

Anxiety and fear are commonly comorbid with depression and recent preclinical work has suggested a protective role for C3a/C3aR in chronic-stress induced depressive-behaviour^91^. Given the common co-occurrence of anxiety and depression, our findings of enhanced anxiety in *C3aR*^*-/-*^ mice might seem at odds with the reported resilience of this strain^-^ to depression-related phenotypes^91^. However, the chronic unpredictable-stress paradigm used in these studies is likely to evoke significant inflammation, and therefore the extent to which our data in acutely stressed animals can be compared is questionable. In summary, our findings add significantly to the evidence that perturbations of the complement system, whether reduced complement activation as in the present work or excessive activation as is predicted by *C4* genetic variants^24,92^, have major and dissociable effects on brain and behavioural phenotypes of relevance to core clinical symptoms of psychiatric disease.

## 5. Author’s contributions

The study was designed by LJW, TH, BPM, WPG and LSW. LJW and TH performed behavioural experiments with assistance from NH. Molecular analyses were performed by SAB, EB, MT, ALM, AIB and LJW. Data interpretation were carried out by JH, MJO, JR, WPG, NH, TRH, BPM, LSW and TH. The manuscript was drafted by LJW, TH, WPG and LSW. All authors approved the final manuscript.

## 6. Acknowledgements

The authors thank Professor Craig Gerard and Professor Marina Botto for provision of the *C3aR*^*-/-*^ and *C5*^*-/-*^ strains respectively, and to Rhys Perry, Pat Mason, Helen Read and other staff at Cardiff University BIOSV for their invaluable animal care and husbandry. This work was supported by a Wellcome Trust Integrative Neuroscience PhD Studentship awarded to LJW (099816/Z/12/Z), a Waterloo Foundation Early Career Fellowship awarded to LJW, a Hodge Centre for Neuropsychiatric Immunology Seed Corn and Project grant awarded to LJW and a Wellcome Trust Strategic Award 100202/Z/12/Z (DEFINE) held by the Neuroscience and Mental Health Research Institute at Cardiff University.

## 7. Competing financial interests

The authors declare no competing financial interests.

## 8. Materials and correspondence

All data from this study are available from the corresponding authors upon reasonable request.

## Notes

### Competing Interest Statement

The authors have declared no competing interest.

