## Supplementary Figures for "Complement C3 and C3aR mediate different aspects of emotional behaviours; relevance to risk for psychiatric disorder"

**Westacott *et al***

### Supplementary Figures

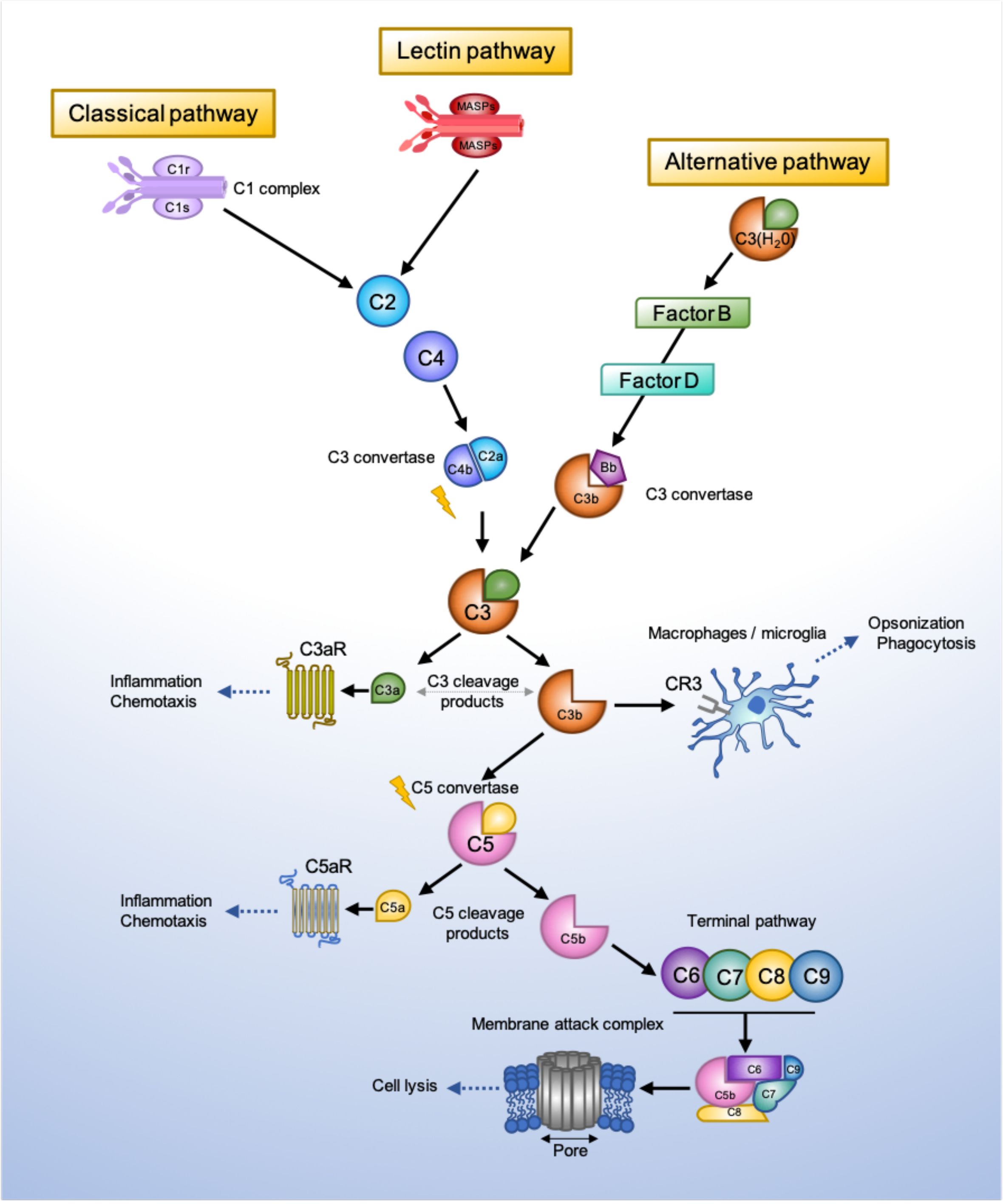


**Figure S1. Overview of the complement system.** Complement system consists of three main 'recognition systems'; the classical pathway, the alternative pathway, the lectin pathway. Each pathway is initiated by different stimuli, which can be either exogenous or endogenous danger signals or pathogen-associated molecular patterns.

Activation of the classical pathway is triggered by detection of antibody-antigen complexes by the initiator molecule of the classical pathway, C1q. C1q is then able to activate C1r and C1s, which cleave C4 and C2 to form the C3 convertase, C4b2a. The lectin pathway is initiated by the recognition of carbohydrates such as mannose upon pathogen surfaces by mannin-binding lectin (MBL), a molecule homologous to C1q of the classical pathway. MBL activates two serine proteases, MASP-1 and MASP-2, which then act to cleave classical pathway components C4 and C2. Activation then proceeds in the same manner as the classical pathway, eventually leading to generation of the C4b2a convertase. The alternative pathway differs from the classical and lectin pathways in that it does not require pathogen recognition to initiate activation. Rather, a continuous state of low-level alternative pathway activation occurs, termed C3 tickover. In this process, the internal thioester bond of circulating C3 molecules is hydrolysed due to nucleophillic attack by $H_20$. This process occurs at a slow yet constant rate, leading to the formation of C3($H_20$). After a series of interactions with C3b and regulatory molecules (Factor B and Factor D), the alternative pathway convertase is formed, C3bBb. Convertase complexes resulting from these distinct pathways then cleave C3, generating the main effectors of the complement system; C3b and C3a. When deposited on surfaces of pathogens or damaged host cells, intact C3b serves to attract phagocytic macrophages, a process known as opsonisation, via complement receptor 3 (CR3). Unbound C3b molecules can also associate with other complement molecules to form a C5 convertase complex. C5 is then cleaved in a similar manner to C3, thereby generating cleavage fragments C5a and C5b. This cascade of activation ultimately leads to assembly of the terminal complement effector; the membrane attack complex (MAC). Aggregation of MAC molecules on a target cell or pathogen creates pores in the cell membrane, leading to death by osmotic cell lysis. The anaphylatoxins generated by C3 and C5 cleavage, C3a and C5a respectively, possess the most potent inflammatory effects of the complement cascade and exert effects within the picomolar to nanomolar range in close proximity to cells bearing the relevant receptors, C3aR and C5aR.

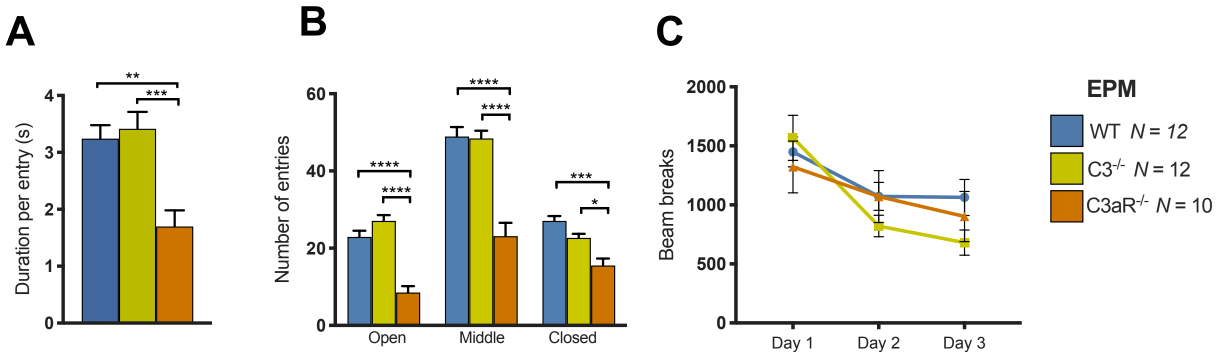


**Figure S2. Supplementary data from the elevated plus maze (EPM) in male mice**

**(A)** *C3aR*$^{-/-}$ mice spent significantly less time (1.70±0.28s) on the open arms per entry than wildtype (3.24±0.23s) and *C3*$^{-/-}$ mice (3.41±0.29s, $F_{2,31}$=11.1, p=0.0002). **(B)** *C3aR*$^{-/-}$ mice were less active in the EPM than wildtype and *C3*$^{-/-}$ mice, making significantly fewer entries to all of the zones of the maze (main effect of GENOTYPE, $F_{2,31}$=29.4, p<0.0001. Open *C3aR*$^{-/-}$ 8.50±1.54 vs. wildtype 22.92±1.63, *C3*$^{-/-}$ 27.08±1.51; Middle *C3aR*$^{-/-}$23.10±3.18 vs. wildtype 48.92±2.45, *C3*$^{-/-}$ 48.42±1.990; Closed *C3aR*$^{-/-}$ 15.50±1.68 vs. wildtype 27.08±1.26, *C3*$^{-/-}$ 22.67±1.05). **(C)** To demonstrate that the reduced activity in the EPM was task/situation specific, locomotor activity was measured in a separate non-anxiety provoking environment, over three days. There were no differences between wildtype, *C3aR*$^{-/-}$ and *C3*$^{-/-}$ mice in total activity in these sessions (main effect of GENOTYPE, $F_{2,31}$=0.344, p=0.712), and all mice demonstrated normal habituation to the test environment between sessions (main effect of DAY, $F_{2,62}$ =61.9, p<0.001). Data are mean ± S.E.M. *, **, *** and **** represent p≤0.05, p≤0.01, p≤0.001 and p≤0.0001 for *post-hoc* genotype comparisons, respectively.

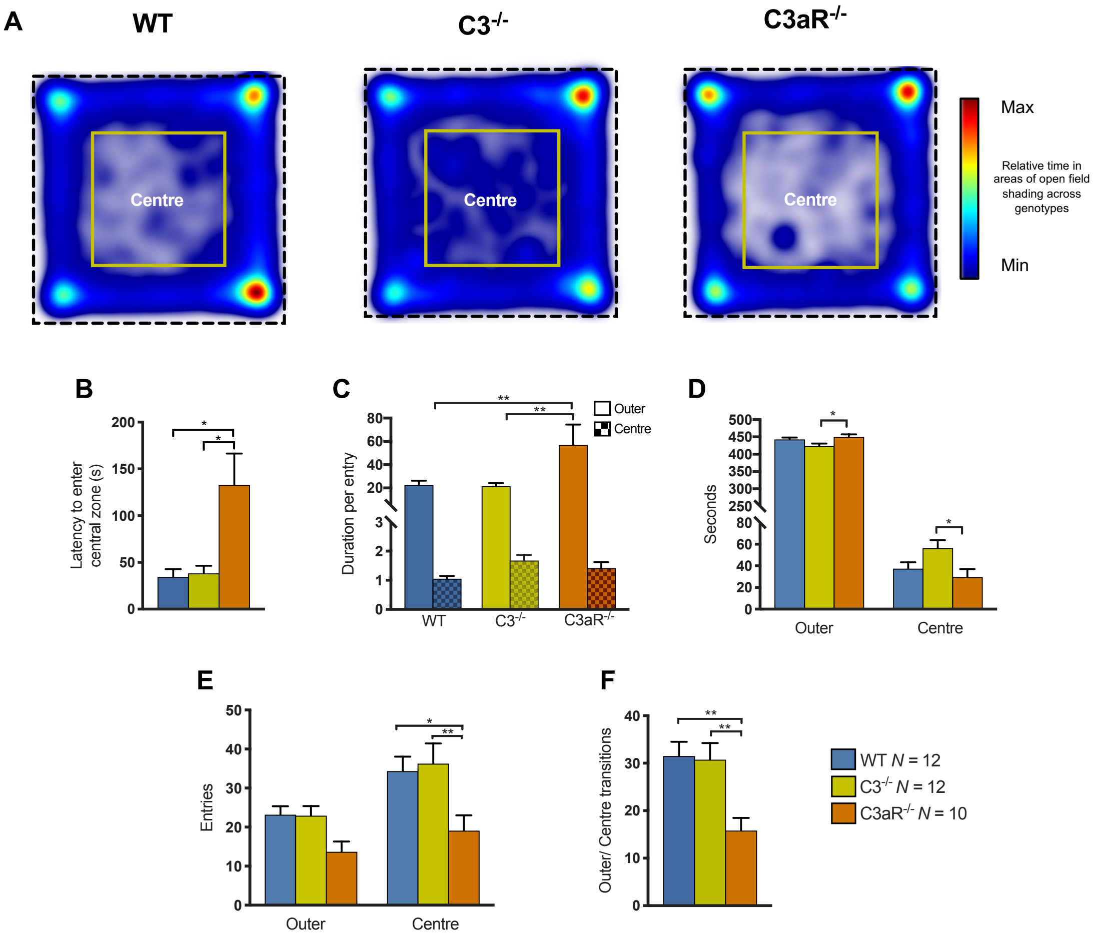


**Figure S3. Male *C3aR*$^{-/-}$ mice displayed anxiety-like behaviour in the open field.** The apparatus consisted of a square-shaped arena (750 x 750 mm, length x width), constructed of white plastic and illuminated evenly at 15 lux. Subjects were placed, facing the centre of one of the walls and allowed to explore the maze for 10 min. The apparatus was subdivided into two virtual concentric zones (shown in **a**) the central region measured 400 * 400 mm$^2$ and the outer zone was defined as regions within 350 mm of each wall. **(A)** Heatmaps indexing relative time in areas (centre and outer) of open field. **(B)** *C3aR*$^{-/-}$ mice took significantly longer (133±33.4s) to initially enter the aversive central zone than wildtype (34.4±8.36s, p=0.0037) and *C3*$^{-/-}$ mice (38.3±8.16s, p=0.119, overall Kruskal-Wallis test $H_2$=12.2, p=0.0022). **(C)** Analysis of duration per entry data revealed a GENOTYPE×ZONE interaction ($F_{2,31}$=4.38, p=0.0211) attributable to *C3aR*$^{-/-}$ mice spending significantly longer in the outer region per entry (wildtype 22.59±3.72s vs *C3aR*$^{-/-}$ 57.03±15.91s p=0.0016, *C3aR*$^{-/-}$ vs *C3*$^{-/-}$ 21.49±2.69s p=0.0011). **(D)** As expected, all subjects displayed thigmotaxis as demonstrated by a significantly greater duration spent in the outer region of the maze (main effect of ZONE, $F_{1,31}$=2541, p<0.0001) and there was a GENOTYPE×ZONE interaction ($F_{2,31}$=4.03, p=0.0279) owing to *C3*$^{-/-}$ subjects spending significantly less time (423.63±7.28s) in the outer region compared to *C3aR*$^{-/-}$ (450.26±6.64s, p=0.0230) mice (Outer; wildtype vs *C3*$^{-/-}$ p=0.1154, wildtype vs *C3aR*$^{-/-}$ p=0.7127). *C3*$^{-/-}$ mice also spent significantly more time (56.37±7.28s) in the central zone compared to *C3aR*$^{-/-}$ mice (29.74±6.64s, p=0.0230) but were not significantly different to wildtype (37.46±5.84s, p=0.1154). **(E)** Due to all subjects spending the majority of the test duration in the outer region, all subjects made a greater number of entries to the centre zone (main effect of ZONE, $F_{1,31}$=56.0, p<0.0001). *C3aR*$^{-/-}$ mice made significantly fewer entries (19.10±3.57) to the central zone than other genotypes however (wildtype 34.33±3.74, *C3*$^{-/-}$ 36.25±5.19, main effect of GENOTYPE, $F_{2,31}$=4.52, p=0.0190). **(F)** *C3aR*$^{-/-}$ mice also made fewer transitions (15.80±2.44) between the central and outer zones than wildtype (31.50±3.02, p=0.0043) and *C3*$^{-/-}$ mice (30.75±3.50, p=0.0066, overall ANOVA $F_{2,31}$=7.44, p=0.0023). Data are mean + S.E.M. * and ** represent p≤0.05 and p≤0.01, for *post-hoc* genotype comparisons, respectively.

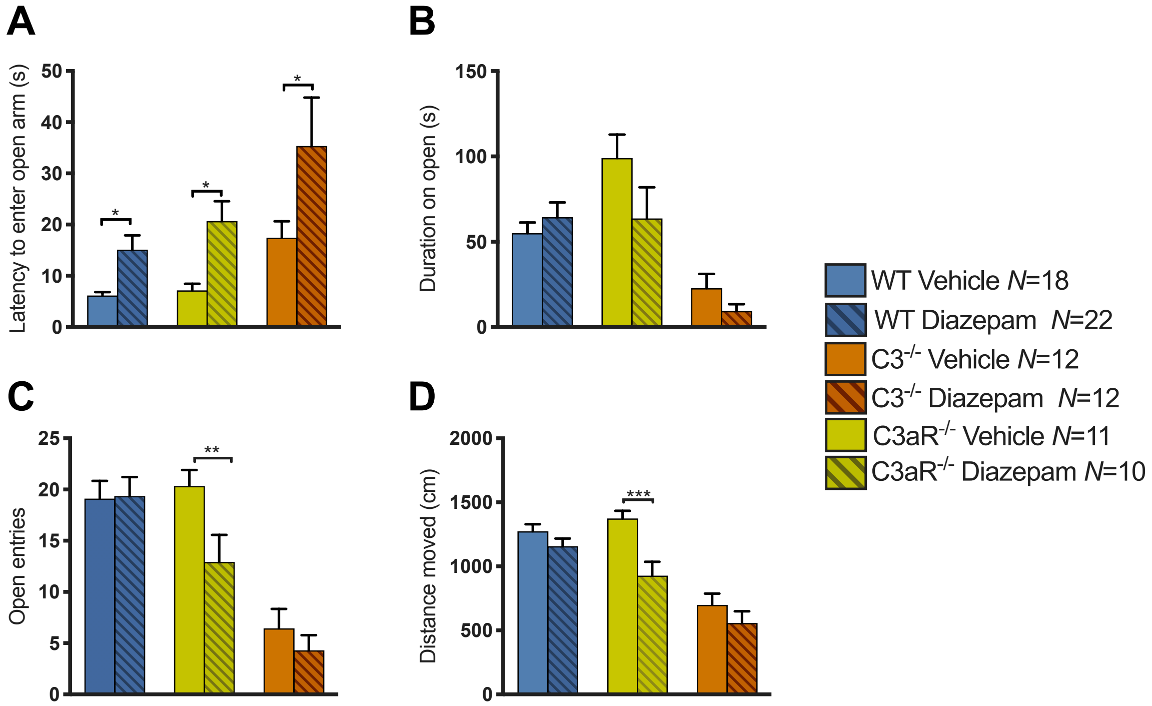


**Figure S4. Altered sensitivity to diazepam in male *C3aR*$^{-/-}$ and *C3*$^{-/-}$ mice. (A)** There was a common effect of diazepam across genotypes whereby the drug significantly increased latency to first open arm visit (28.29±5.50s) compared to vehicle treated animals (13.26±4.43s, main effect of DRUG, $F_{1,73}$=23.6, p<0.0001). As previously, *C3aR*$^{-/-}$ mice took significantly longer (44.61±12.20s) than other groups (wildtype 12.23±2.06s, *C3*$^{-/-}$ 19.40±6.37s) to initially enter the open arm (main effect of GENOTYPE, $F_{2,69}$=10.6, p<0.0001). This was unlikely to be a sedative effect (see distance moved shown in D) **(B)** There was a main effect of GENOTYPE ($F_{2,79}$=15.6, p<0.0001), reflecting reduced open arm duration in *C3aR*$^{-/-}$ mice (16.43±4.93s) compared to wildtype (60.23±5.50s, p=0.0003) and C3-/- mice (81.35±11.79s, p<0.0001). There was no main effect of DRUG ($F_{1,79}$=2.14, p=0.1477) on open arm duration. **(C)** There was a main effect of GENOTYPE ($F_{2,79}$=24.5, p<0.0001) and no main effect of DRUG ($F_{1,69}$=2.14, p=0.1477) on open arm entries. However *post hoc* tests demonstrated that diazepam reduced open arm entries in *C3*$^{-/-}$ mice (vehicle

20.33±1.57 vs. diazepam 12.92±2.66, p=0.496). **(D)** There were also genotype differences in distance moved around the maze (main effect of GENOTYPE, $F_{2,79}$=32.1, p <0.0001) and a GENOTYPE × DRUG interaction approaching significance ($F_{2,79}$=2.85, p =0.0640). Diazepam treatment significantly reduced distance moved in *C3*$^{-/-}$ mice only (*C3*$^{-/-}$ vehicle 1374.22±59.65 vs. *C3*$^{-/-}$ diazepam 925.42±109.01, p=0.0006). Data are mean + S.E.M. *, ** and *** represent p≤0.05, p≤0.01 and p≤0.001, for *post-hoc* genotype comparisons, respectively.

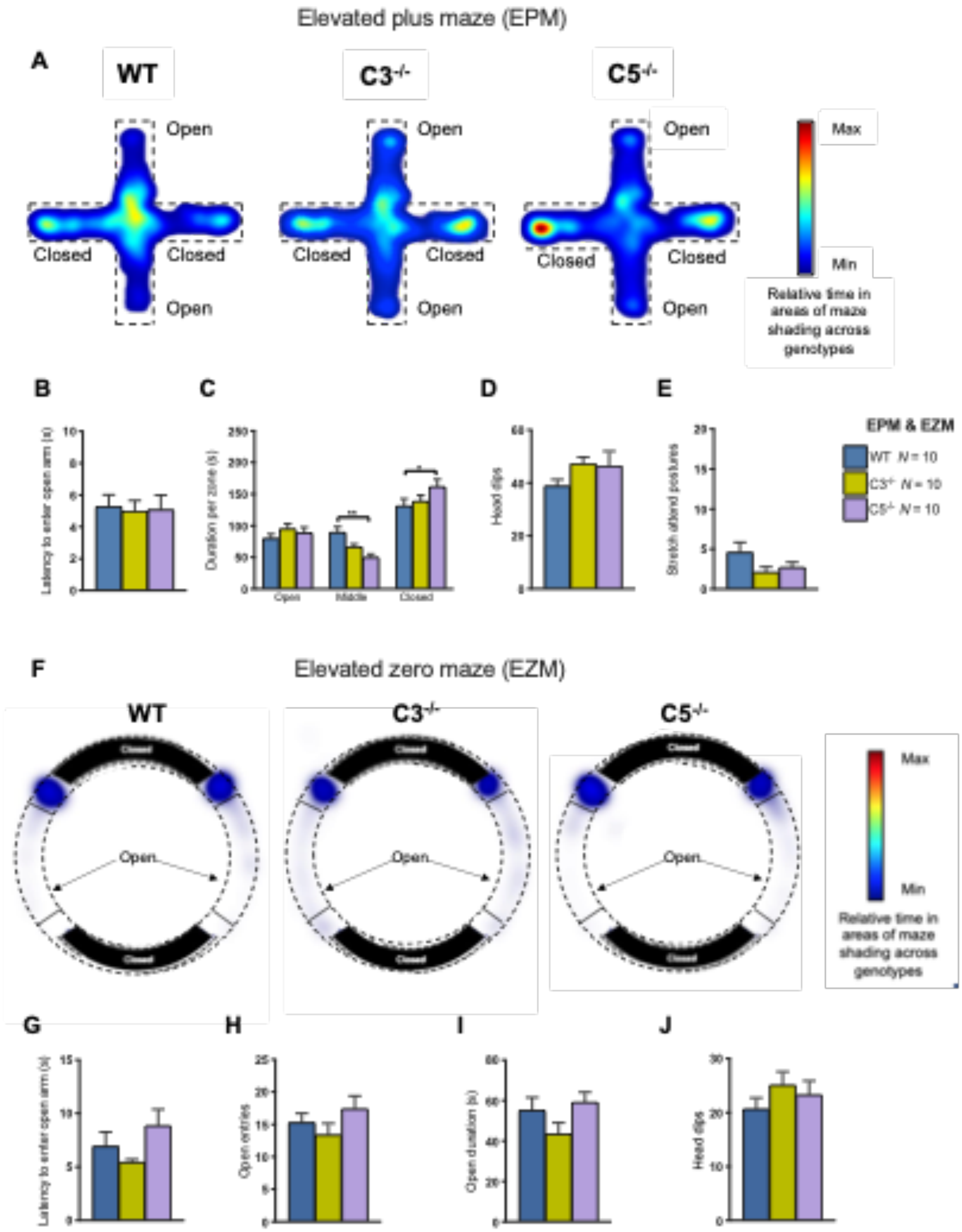


**Figure S5.** ***C5***$^{-/-}$ **mice demonstrated normal reactivity to the elevated plus maze (EPM) and elevated zero maze (EZM) (A)** Heatmaps demonstrating duration spent per zone of the EPM. **(B)** There were no differences in latency to first enter the open arm ($F_{2,27}$=0.372, p= 0.9636) **(C)** Duration (s) spent per zone (open, middle, closed)

of the EPM. There was no main effect of GENOTYPE ($F_{2,27}$=0.183, p=0.8339) but there was a significant main effect of ZONE ($F_{2,54}$=41.2, p<0.0001) and a significant GENOTYPE×ZONE interaction ($F_{4,54}$=3.18, p=0.0203). Whilst there were no differences in time spent on the open arm, *C5*$^{-/-}$ mice spent significantly less time in the middle zone (50.3±4.99 s) than wildtype (89.40±8.97 s, p=0.0049), and more time in the closed zone (161.86±11.56 s) than wildtype mice (131.12±11.45 s, p=0.0356). There were no significant differences in the number of **(D)** head dips (Kruskal-Wallis test $H_{2,29}$=4.37, p=0.1126) or **(E)** stretch attend postures ($F_{2,27}$=2.12, p=0.1401) **(F)** Heatmaps demonstrating duration spent per zone of the EZM. There were no significant differences between genotypes in latency to first open arm entry **(G)** ($F_{2,27}$=2.13, p=0.1388), total open arm entries **(H)** ($F_{2,27}$=1.34, p=0.2780), duration spent on the open arms **(I)** ($F_{2,27}$=2.11, p=0.1411) or head dips **(J)** ($F_{2,27}$=0.855, p=0.4364). Data are mean + S.E.M. * and ** represent p≤0.05 and p≤0.01 for *post-hoc* genotype comparisons, respectively.

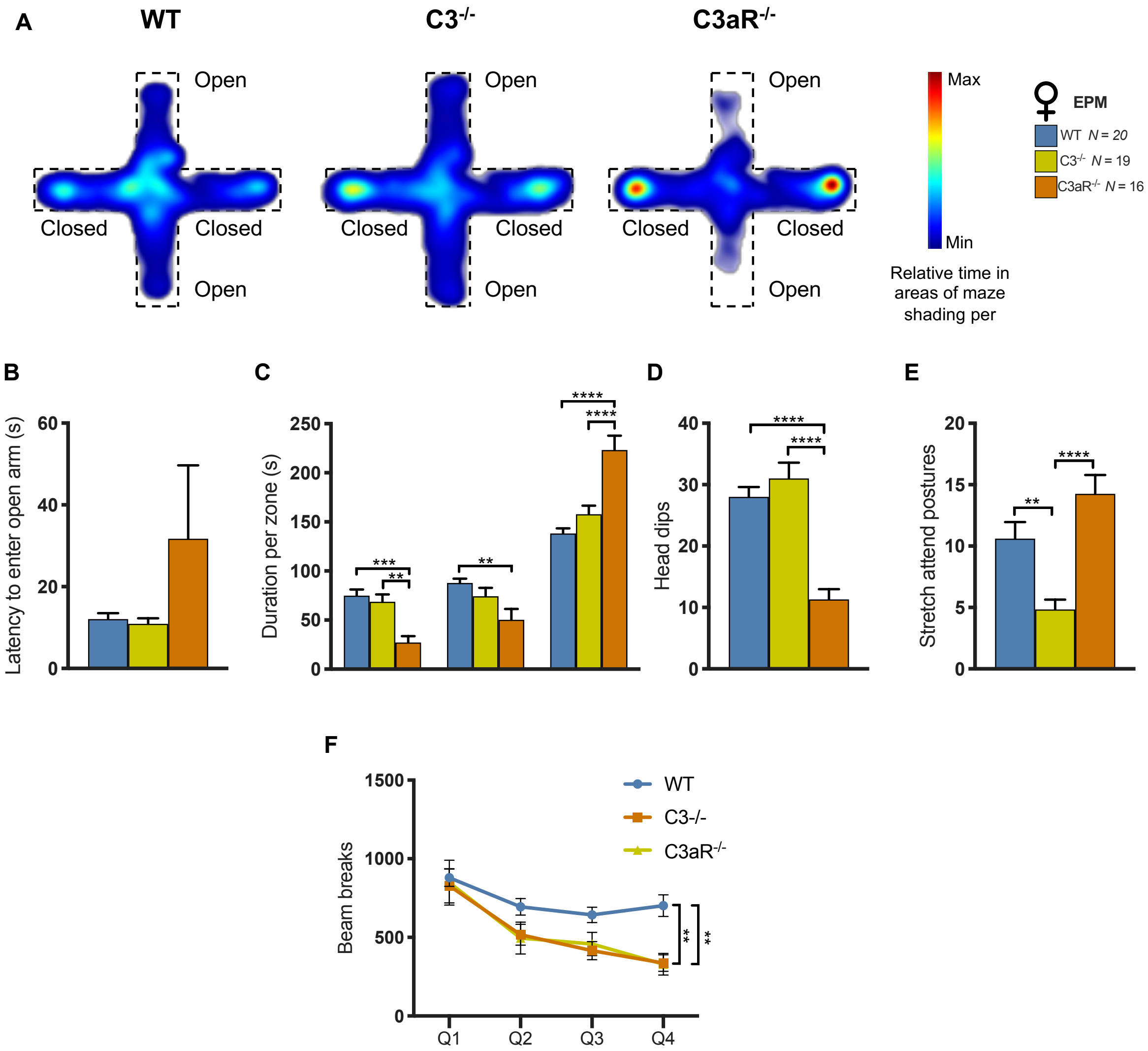


**Figure S6. Female *C3aR*$^{-/-}$ mice also show increased anxiety-like behaviour in the elevated plus maze (A)** Heatmaps displaying relative time per zone of the EPM across genotypes **(B)** There was a trend towards increased latency to enter the open arm in *C3aR*$^{-/-}$ mice (31.7±17.9s) compared to wildtype (12.1±1.48) and *C3*$^{-/-}$ mice (10.9±1.39s) however this did not reach statistical significance ($H_2$=4.19, p=0.1231) **(C)** *C3aR*$^{-/-}$ mice distributed their time across the EPM differently to wildtype and *C3*$^{-/-}$ mice (GENOTYPE×ZONE, $F_{4,104}$=13.9, p<0.0001), spending less time in the open arms (*C3aR*$^{-/-}$ 26.28±5.66s vs. wildtype 74.93±5.63s, p=0.0003, *C3aR*$^{-/-}$ vs. *C3*$^{-/-}$ 68.75±6.23s p=0.0020) and significantly more time in the closed arms (*C3aR*$^{-/-}$ 225.37±12.79s vs. wildtype 138.26±4.65s, p<0.0001, and *C3*$^{-/-}$ 157.71±7.48s p<0.0001). **(D)** *C3aR*$^{-/-}$ (11.3±1.67) mice performed significantly fewer head dips than wildtype (28.0±1.62, p<0.0001) and *C3*$^{-/-}$ mice (31.0±2.60, p<0.0001, overall ANOVA $F_{2,52}$=25.1, p<0.0001). **(E)** *C3aR*$^{-/-}$ mice performed significantly more stretch attend postures (SAPs; 14.3±1.54) than wildtype (10.6±1.36, p=0.0042) and *C3*$^{-/-}$ mice (4.84±0.79, p<0.0001; overall ANOVA $F_{2,52}$=13.9, p<0.0001). **(F)** Prior to testing on the EPM, locomotor activity was assessed in a non-anxiety provoking environment. Animals were placed into activity boxes for 120 minutes and their locomotion was recorded and analysed in four 30 minute-quartiles. There was a significant GENOTYPE×TIME interaction ($F_{6,171}$=4.24, p<0.0005) whereby *C3*$^{-/-}$ and *C3aR*$^{-/-}$ mice made significantly fewer beam breaks in the fourth quartile of the test compared to wildtype (wildtype 702.21±69.32, *C3*$^{-/-}$ 336.90±52.82, *C3aR*$^{-/-}$ 329.00±69.70, wildtype vs. *C3*$^{-/-}$ p=0.0011, wildtype vs. *C3aR*$^{-/-}$ p=0.0019, *C3*$^{-/-}$ vs. *C3aR*$^{-/-}$ p=0.9973). Data are mean ± S.E.M. *, **, *** and **** represent p≤0.05, p≤0.01, p≤0.001 and p≤0.0001 for *post-hoc* genotype comparisons, respectively.

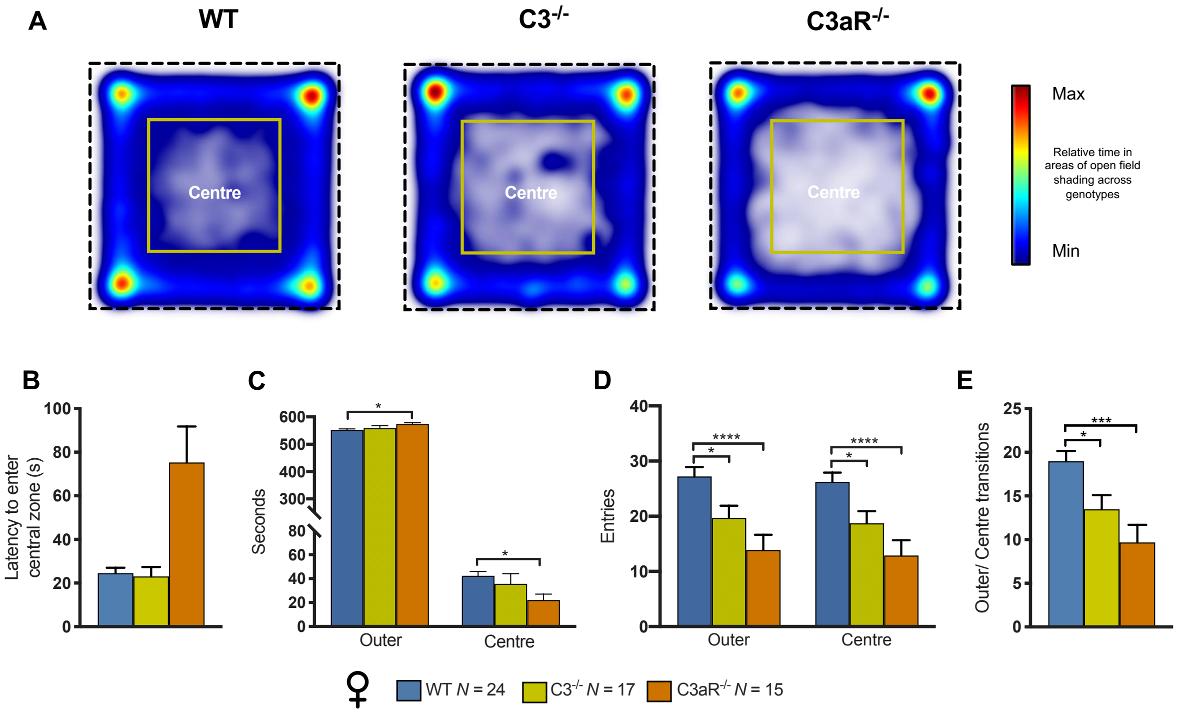


**Figure S7. Female *C3aR*$^{-/-}$ mice also displayed anxiety-like behaviour in the open field. (A)** Heatmaps indexing relative time in areas (centre and outer) of open field. **(B)** There was a trend indicative of *C3aR*$^{-/-}$ taking significantly longer (75.3±16.5s) to initially enter the aversive central zone than wildtype (24.6±2.47s) and *C3*$^{-/-}$ mice (23.1±4.23s) that was approaching significance (Kruskal-Wallis test $H_2$=5.44, p=0.0658). **(C)** As expected, subjects displayed thigmotaxis as demonstrated by a significantly greater duration spent in the outer region of the maze (main effect of ZONE, $F_{1,53}$=6092 p<0.0001) and there was a GENOTYPE×ZONE interaction ($F_{2,53}$=3.21, p=0.0484) owing to *C3aR*$^{-/-}$ subjects spending significantly more time (573.65±4.93s) in the outer region than wildtype mice (552.72±3.86s, p=0.0318) and less time in the central region (22.05±4.91s) than wildtype mice (42.41±3.65s, p=0.0380). There were no significant differences between *C3aR*$^{-/-}$ and *C3*$^{-/-}$ mice in the outer (562.56±7.58s, p=0.2265) or central zones (31.59±7.53s, p=0.2720). **(E)** Due to all subjects spending the majority of the test duration in the outer region, all subjects made a greater number of entries to the centre zone (main effect of ZONE, $F_{1,53}$=938,

$p<0.0001$). There was also a significant main effect of GENOTYPE ($F_{2,53}=10.1$, $p=0.0002$) whereby *C3aR*$^{-/-}$ mice made significantly fewer entries overall (13.62±2.75) compared to other genotypes (wildtype 26.72±1.81, *C3*$^{-/-}$ 19.20±2.19). **(F)** Both *C3*$^{-/-}$ (13.5±1.62) and *C3aR*$^{-/-}$ (9.67±2.02) mice made fewer transitions between the central and outer zones than wildtype (19.0±1.20, vs. *C3*$^{-/-}$ $p=0.0327$, vs. *C3aR*$^{-/-}$ $p=0.0003$, overall ANOVA $F_{2,53}=9.39$, $p=0.0003$). Data are mean + S.E.M. * and ** represent $p\leq0.05$ and $p\leq0.01$, for *post-hoc* genotype comparisons, respectively.

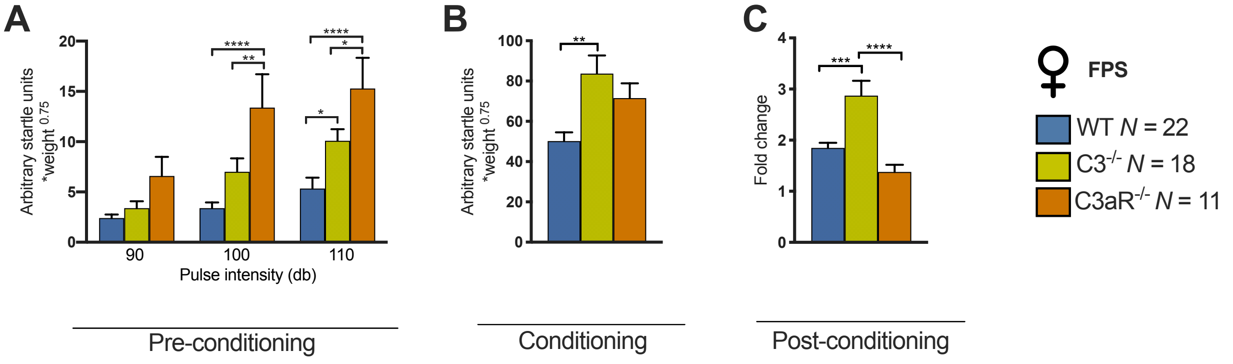


**Figure S8. Female *C3*$^{-/-}$ but not *C3aR*$^{-/-}$ mice also demonstrated enhanced fear-potentiated startle. (A)** In the pre-conditioning session, there was a significant main effect of GENOTYPE ($F_{2,48}$=11.3, p<0.0001), STIMULUS INTENSITY ($F_{2,96}$=30.2, p<0.0001) and a significant GENOTYPE × STIMULUS INTENSITY interaction ($F_{4,96}$=3.11, p=0.0187). *C3aR*$^{-/-}$ mice demonstrated increased levels of startle responding relative to wildtype mice at 100dB (*C3aR*$^{-/-}$ 13.39±2.94 vs. wildtype 3.40±0.54, p<0.0001, *C3aR*$^{-/-}$ vs. *C3*$^{-/-}$ 7.02±1.25, p=0.0089) where as both *C3*$^{-/-}$ and *C3aR*$^{-/-}$ mice demonstrated greater startle responses at 110dB (*C3*$^{-/-}$ 10.10±1.09 vs. wildtype 5.35±1.03, p=0.0217, *C3aR*$^{-/-}$ 15.29±2.70 vs. wildtype p<0.0001, *C3*$^{-/-}$ vs. *C3aR*$^{-/-}$ p=0.0418). **(B)** *C3*$^{-/-}$ mice showed increased startle responses to the footshock+CS pairings (83.72±8.56) relative to wildtype mice (50.20±4.17, p=0.0017, overall ANOVA $F_{2,48}$=6.97, p=0.0022). **(C)** In the post-conditioning session, all mice demonstrated increases to the pulse+CS stimuli in comparison to pulse-alone stimuli, as demonstrated by the fold-change increase in startle responding, however, this effect was significantly increased in *C3*$^{-/-}$ mice (3.18±0.25) relative to wildtype (1.96±0.09, p=0.0072) and *C3aR*$^{-/-}$ mice (1.58±0.10, p<0.0001, overall Kruskal-Wallis test $H_2$=21.1, p<0.0001). Data are mean + S.E.M. *, **, *** and **** represent p≤0.05, p≤0.01, p≤0.001 and p≤0.0001 for *post-hoc* genotype comparisons, respectively.

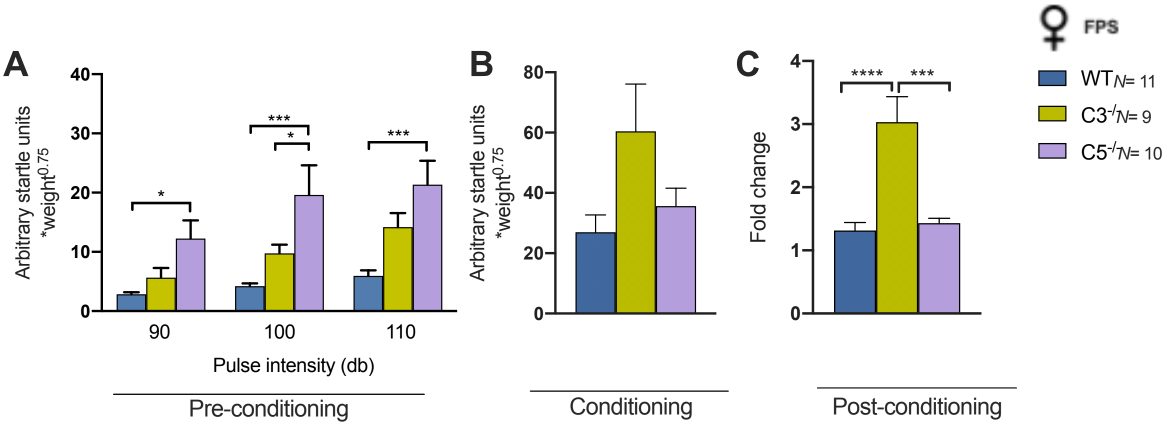


**Figure S9. Female *C5*$^{-/-}$ mice demonstrated normal fear-potentiated startle responses. (A)** In the pre-conditioning session, there was a significant main effect of GENOTYPE ($F_{2,27}$=9.59, p=0.0007) and STIMULUS INTENSITY ($F_{2,54}$=13.8, p<0.0001). The GENOTYPE × STIMULUS INTENSITY interaction term was not significant ($F_{4,54}$=1.47, p=0.2227). *C5*$^{-/-}$ mice demonstrated increased levels of startle responding relative to wildtype mice at 90dB (*C5*$^{-/-}$ 12.26±3.07 vs. wildtype 2.85±0.32, p=0.0270) and 110dB (*C5*$^{-/-}$ 21.37±4.03 vs. wildtype 5.94±0.94, p=0.0001). At 100dB, *C5*$^{-/-}$ mice (19.64±4.98) had significantly higher startle responses than both wildtype (4.22±0.47, p=0.0001) and *C3*$^{-/-}$ mice (9.75±1.49, p=0.0273). **(B)** *C3*$^{-/-}$ mice showed a trend towards increased startle responses to the footshock+CS pairings (60.44±15.62) relative to wildtype (26.98±5.70) and *C5*$^{-/-}$ mice (35.63±5.90) but this difference did not reach significance (Kruskal-Wallis test $H_2$=3.60, p=0.1656). **(C)** In the post-conditioning session, all mice demonstrated the expected potentiation of startle response to pulse+CS trials in comparison to pulse-alone trials. Again, this effect was significantly increased in *C3*$^{-/-}$ mice (3.03±0.40) relative to wildtype (1.31±0.13, p<0.0001) and *C5*$^{-/-}$ mice (1.43±0.08, p=0.0001, overall one-way ANOVA $F_{2,27}$=16.8, p<0.0001). *C5*$^{-/-}$ mice were not significantly different to wildtypes (p=0.9251). Data are mean + S.E.M. *, *** and **** represent $p \leq 0.05$, $p \leq 0.001$ and $p \leq 0.0001$ for *post-hoc* genotype comparisons, respectively.
